# Plectin Ensures Intestinal Epithelial Integrity and Protects Colon Against Colitis

**DOI:** 10.1101/2020.10.06.323493

**Authors:** Alzbeta Krausova, Petra Buresova, Lenka Sarnova, Gizem Oyman-Eyrilmez, Jozef Skarda, Pavel Wohl, Lukas Bajer, Eva Sticova, Lenka Bartonova, Jiri Pacha, Gizela Koubkova, Jan Prochazka, Marina Spörrer, Christopher Dürrbeck, Zuzana Stehlikova, Martin Vit, Natalia Ziolkowska, Radislav Sedlacek, Daniel Jirak, Miloslav Kverka, Gerhard Wiche, Ben Fabry, Vladimir Korinek, Martin Gregor

## Abstract

Plectin, a highly versatile cytolinker protein, provides tissues with mechanical stability through the integration of intermediate filaments (IFs) with cell junctions. Here, we hypothesize that plectin-controlled cytoarchitecture is a critical determinant of the intestinal barrier function and homeostasis. Mice lacking plectin in intestinal epithelial cells (IEC; *Ple^ΔIEC^*) spontaneously developed colitis characterized by extensive detachment of IECs from the basement membrane (BM), increased intestinal permeability, and inflammatory lesions. Moreover, plectin expression was reduced in colons of ulcerative colitis (UC) patients and negatively correlated with the severity of colitis. Mechanistically, plectin deficiency in IECs led to aberrant keratin filament (KF) network organization and formation of dysfunctional hemidesmosomes (HDs) and intercellular junctions. In addition, the hemidesmosomal α6β4 integrin (Itg) receptor showed attenuated association with KFs, and protein profiling revealed prominent downregulation of junctional constituents. Consistent with effects of plectin loss in the intestinal epithelium, plectin-deficient IECs exhibited remarkably reduced mechanical stability and limited adhesion capacity *in vitro*. Feeding mice with a low-residue liquid diet that reduced mechanical stress and antibiotic treatment successfully mitigated epithelial damage in the *Ple^ΔIEC^* colon.

## INTRODUCTION

The intestinal epithelium is composed of a single layer of tightly linked IECs, forming a selective physical barrier that is critical for gut homeostasis. A breach in the intestinal barrier, referred to as “leaky gut”^1^, results in excessive exposure to luminal microbiota and in a concomitant innate immune response. Subsequent dysregulation of finely-tuned interplay among gut microbiota, IECs, and immune cells accounts for uncontrolled inflammation and pathogenesis of intestinal disorders such as inflammatory bowel disease (IBD) and colorectal cancer (CRC)^2, 3^.

The epithelial barrier function is secured by cell junctions that seal intercellular spaces and interlink IECs with the underlying BM into a structural and functional continuum. Alterations in junctional proteins and BM components may lead to a breakdown of the barrier, and genetic studies identified multiple links between junction/BM-associated genes and the development of IBD^4–6^. While apical tight junctions (TJs) and subjacent adherens junction (AJs) confer paracellular transport selectivity, desmosomes (Ds) together with BM-linked hemidesmosomes (HDs) provide the intestinal epithelium with resilience to mechanical stress generated by intestinal peristalsis^7^. It is noteworthy that recently reported mouse models demonstrate the protective role of Ds and HDs in the context of both intestinal inflammation^8, 9^ and colitis-associated CRC^8^. Accumulating evidence suggests that fundamental features of Ds and HDs (such as stability, dynamics, and mechanotransduction capacity) heavily rely on their interconnection with KF networks^10, 11^. This places plakins^12^, a family of cytolinker proteins mediating physical linkage between KFs and cell junctions, at the very center of the processes controlling epithelial homeostasis.

Plectin, a highly versatile member of the plakin protein family, crosslinks intermediate filaments (IFs) of different types and anchors them at cellular junctions, including HDs and Ds of epithelial cells^13^. Multiple mutations in the *plectin* gene have been identified in epidermolysis bullosa (EB)^14^, a disorder characterized by excessive blister formation in skin^15, 16^ with reported cases of concurrent IBD^17, 18^. Previous studies have shown that plectin ablation disrupts highly organized epithelial KF networks and alters the structure and functionality of cell junctions^19–22^. For example, a tissue-specific deletion of *plectin* in the mouse biliary epithelium has adverse effects on the formation of TJs, AJs, and Ds, with deleterious consequences for epithelial stability under cholestasis^22^. Likewise, analysis of knock-in mice recapitulating dominant EB simplex suggests that HD stability in basal keratinocytes depends on plectin-mediated recruitment of KFs^20^. Mechanistically, dysfunctional HDs account for epithelial fragility and lesional defects^23^ which resemble those seen in patients with IBD^24^. Although these observations suggest a linkage between plectin dysfunction and intestinal pathologies, plectin’s role in the intestinal epithelium remains unaddressed.

In this study, we found that plectin expression was reduced in patients with UC, and that plectin expression levels negatively correlated with the severity of colitis. To study the underlying molecular mechanisms, we generated two new mouse lines: one constitutive (*Ple^ΔIEC^*) and the other with tamoxifen (TMX)-inducible (*Ple^ΔIEC-ERT2^*) plectin ablation in IECs. The phenotypic characterization of these mice demonstrated that loss of plectin leads to spontaneous development of a colitic phenotype characterized by extensive detachment of IECs from the BM, increased intestinal permeability, and formation of inflammatory lesions. These results demonstrate absolute indispensability of plectin for the maintenance of intestinal epithelium integrity, and moreover that both mouse lines provide a useful model systems for investigating disease etiology and testing palliative therapies.

## RESULTS

### Suppression of plectin in human patients with UC

To examine the role of plectin in the pathogenesis of UC, we screened for potential alterations of plectin expression in a cohort of more than 50 UC patients. The analysis of immunolabelled biopsy samples taken from patients and healthy controls revealed discontinuous and rather patchy plectin staining patterns in UC biopsies. In healthy controls, plectin decorated both apical and basal membranes of IECs evenly (Figure 1A), resembling plectin localization in mouse intestinal sections^25^, and Figure S1A). In addition, mRNA profiling showed significantly reduced expression levels of plectin in UC biopsies (Figure 1B). Histological analysis revealed that low mRNA levels of plectin were associated with higher inflammation (Figure 1C) and higher C reactive protein levels in serum (not shown).

**Figure 1.**
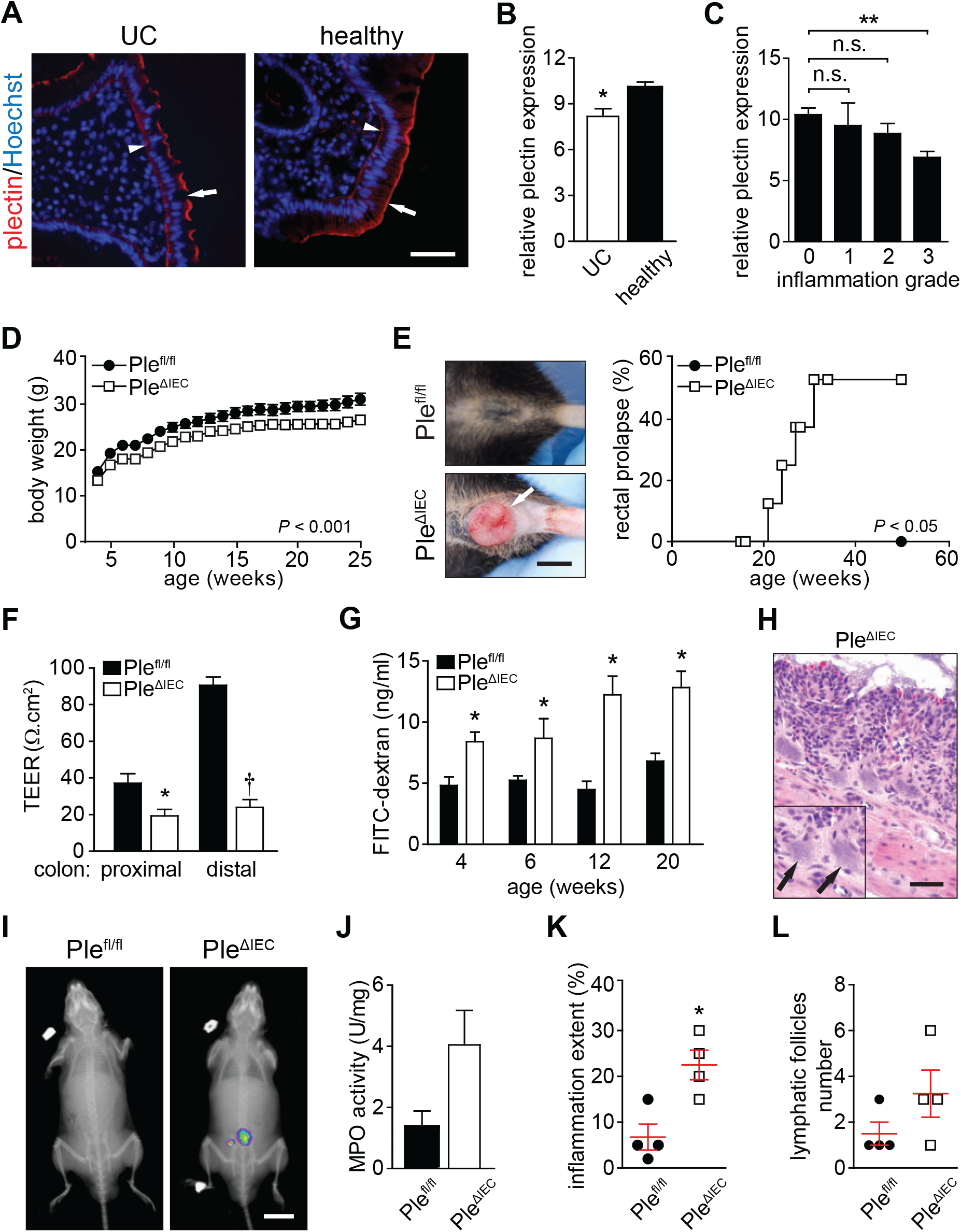
Loss of plectin is associated with UC in human patients and leads to intestinal epithelial barrier dysfunction with concomitant inflammation in mouse. (A) Paraffin-embedded colon sections from healthy controls (healthy) and UC patients (UC) were immunolabelled with antibodies to plectin (red). Nuclei were stained with Hoechst (blue). Arrows, apical IEC membrane; arrowheads, basal IEC membrane. Scale bar, 50 μm. (B) Relative *plectin* mRNA levels in rectum biopsies collected from healthy controls and UC patients. n = 18-51. (C) Relative *plectin* mRNA expression in rectum biopsies collected from UC patients clustered based on inflammation scored in H&E-stained rectum sections. n = 6-30. (D) Body weight of *Ple^fl/fl^* and *Ple^ΔIEC^* mice was monitored for 25 weeks. n = 7. (E) Representative images of rectum of 30-week-old *Ple^fl/fl^* and *Ple^ΔIEC^* mice. Kaplan-Meier graph shows age-related rectal prolapse incidence. (F) Intestinal transepithelial electrical resistance (TEER) measured *ex vivo* in both proximal and distal colons of *Ple^fl/fl^* and *Ple^ΔIEC^* mice. n = 4. (G) *In vivo* permeability of mucosa of *Ple^fl/fl^* and *Ple^ΔIEC^* mice (at age indicated) measured by monitoring 40-kDa FITC-dextran levels in plasma 4 h after orogastric gavage. n = 3-7. (H) Representative image of *Ple^ΔIEC^* colon section stained with H&E. Arrows, bacterial patches in mucosa. Scale bar, 50 μm. (I) *In vivo* chemiluminescence images of *Ple^ΔIEC^* and *Ple^fl/fl^* mice injected with myeloperoxidase (MPO) inflammation probe. (J) MPO activity (marker of neutrophil infiltration) measured in colon lysates from *Ple^ΔIEC^* and *Ple^fl/fl^* mice. n = 3. (K,L) Inflammation extent (percentage) (K) and the number of lymphatic follicles (L) assessed from H&E-stained sections of *Ple^fl/fl^* and *Ple^ΔIEC^* colons. n = 4. Data are presented as mean ± SEM, \**P* < 0.05, \*\**P* < 0.01, †*P* < 0.001.

### IEC-specific plectin-deficient mice develop a colitic phenotype due to intestinal barrier dysfunction

To explore the role of plectin in the intestinal epithelium in greater detail, we generated IEC-specific *plectin* knockout (*Ple^ΔIEC^*) mice. Successful ablation of plectin in IECs was confirmed by immunofluorescence microscopy (Figure S1B). The newly generated *Ple^ΔIEC^* mice had a considerably lower body weight (Figure 1D and Figure S1C), suffered from persistent diarrhea with occasional rectal bleeding (Figure S1D, and not shown), and frequently developed rectal prolapse (Figure 1E). As the onset and progression of UC correlate with defects in the intestinal barrier function^26, 27^, we assessed barrier integrity either by *ex vivo* measurements of intestinal transepithelial electrical resistance (TEER) or by *in vivo* orogastric gavage of FITC-dextran. We observed significantly lower TEER in the proximal and distal colon regions of 12-week-old *Ple^ΔIEC^* compared to *Ple^fl/fl^* mice. Moreover, TEER values in *Ple^fl/fl^* mice were three-times higher in their distal parts than in their proximal parts; by contrast, TEER values in *Ple^ΔIEC^* mice were equally low in distal and proximal colon segments (Figure 1F). Compared to *Ple^fl/fl^* mice, *Ple^ΔIEC^* mice consistently displayed a higher penetration rate of FITC-dextran into blood already at 4 weeks, and this difference became even more apparent in older animals (Figure 1G).

Additionally, histological inspection of hematoxylin-eosin (H&E)-stained colon sections revealed extensive translocation of luminal bacteria into *Ple^ΔIEC^* mucosa (Figure 1H). Given the hampered barrier function in the *Ple^ΔIEC^* intestine, we screened *Ple^fl/fl^* and *Ple^ΔIEC^* mice for signs of inflammation. Chemiluminescence-based whole body imaging^28^ showed positive abdominal areas in *Ple^ΔIEC^* mice (Figure 1I), which correlated strongly with significantly higher myeloperoxidase activity (MPO; Figure 1J). Moreover, mild inflammation of the *Ple^ΔIEC^* colon was confirmed by increased immune cell infiltration, extent (or intensity) of acute/chronic inflammation, and lymphatic follicle size (Figure 1K, L and Figure S2A), and a higher percentage of edema and ulceration indicated higher epithelial damage (Figure S2B). Together, these results suggest that plectin is critical for the maintenance of the intestinal barrier and thus could be directly linked to the onset and progression of UC.

### Loss of plectin leads to hyperproliferation and aberrant differentiation of IECs

Further histological inspection of H&E-stained colonic sections revealed thickening of the colonic mucosa and significant crypt damage with excessive sloughing of IECs detached from the subjacent BM in plectin-deficient specimens (Figure 2A). In addition, the colon of *Ple^ΔIEC^* mice showed a higher rate of proliferation as determined from Ki-67-stained sections (Figure 2B). Consistently, an increase in the number of proliferating transit-amplifying IECs in the crypts of *Ple^ΔIEC^* animals was evident from BrdU incorporation assessed 2, 24, and 48 hours after a BrdU pulse (Figure 2C). Interestingly, TUNEL staining indicated a minimal degree of spontaneous apoptosis in both *Ple^fl/fl^* and *Ple^ΔIEC^* mice (Figure S3A). In parallel with the prominent hyperplasia, the *Ple^ΔIEC^* colon contained a higher proportion of PAS-positive goblet cells (Figure 2D), corresponding to a higher mucus discharge (Figure 2E). Immunolabelled *Ple^ΔIEC^* colonic sections also showed lower percentage of chromogranin A (ChgA)-positive enteroendocrine cells (Figure S3B) and an extended keratin 20 (K20)-positive zone (Figure S3C). Similar, albeit less pronounced, trends were observed in the *Ple^ΔIEC^* small intestine (Figure S4). Plectin deficiency thus results in hyperproliferation and aberrant differentiation of IECs, affecting the spatiotemporal organization of the intestinal epithelium.

**Figure 2.**
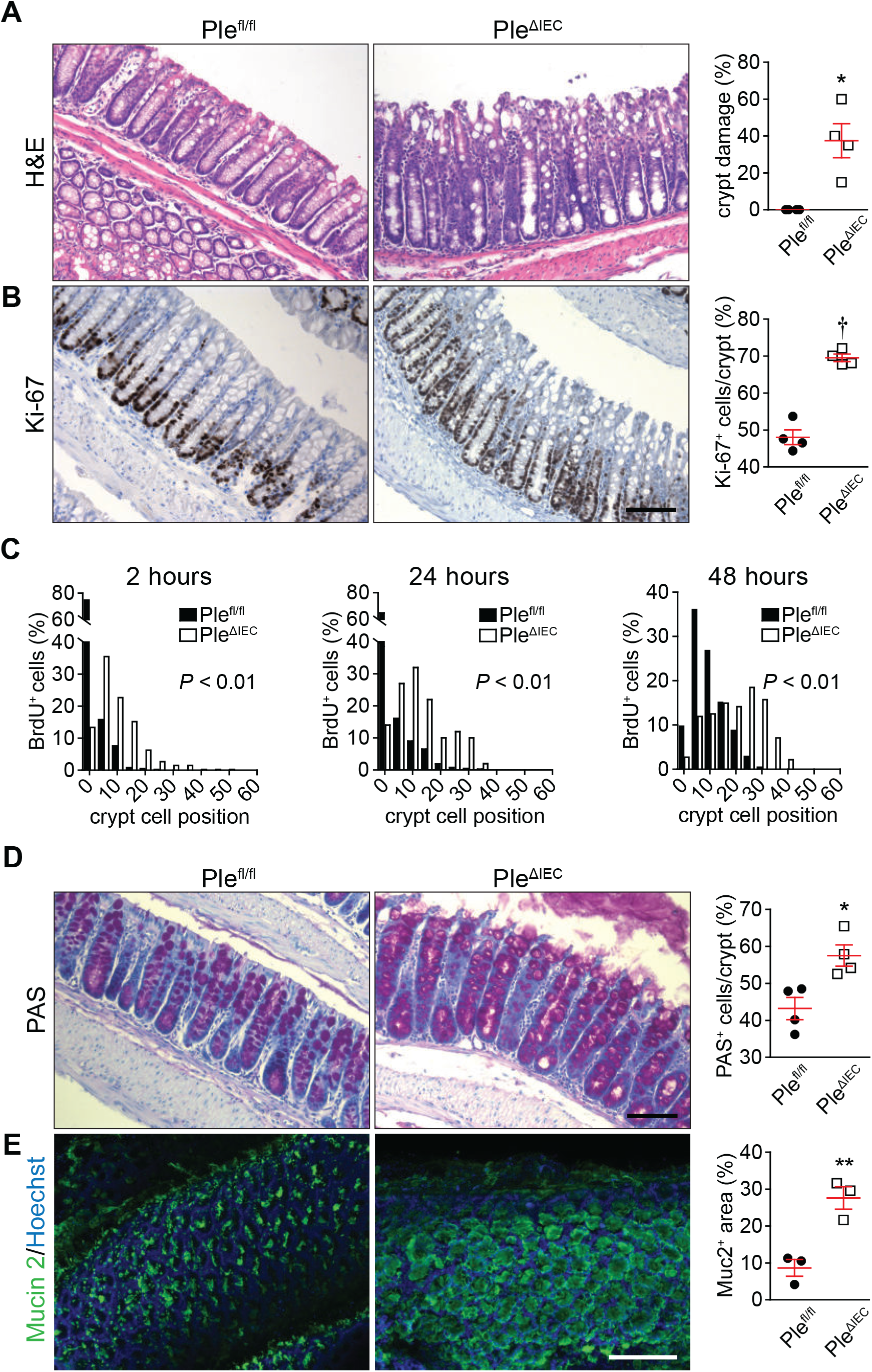
Plectin-deficient IECs exhibit aberrant proliferation and differentiation, resulting in altered crypt organization. (A,B) Representative images of H&E staining (A) and Ki-67 immunohistochemistry (proliferating cells) (B) of *Ple^fl/fl^* and *Ple^ΔIEC^* paraffin-embedded colon sections. Scale bar, 100 μm. Graphs show quantification of colonic crypt damage given as percentage of crypts with >5% of IECs detached from BM (A) and percentage of the Ki-67-positive (Ki-67^+^) IECs per crypt (B). n = 3-4. (C) Histograms showing percentage of BrdU-positive (BrdU^+^) cells in given positions of *Ple^fl/fl^* and *Ple^ΔIEC^* colonic crypts at 2, 24, and 48 hours after BrdU pulse. Cells were numbered sequentially from crypt base to lumen, with cell position 0 assigned to the first cell at the base of each crypt. At least 9 crypts per mouse were analyzed from 3 mice per time point and genotype. (D,E) Representative images of PAS staining (goblet cells) (D) and Mucin 2 (Muc2) immunofluorescence in mucus layer (E) of *Ple^fl/fl^* and *Ple^ΔIEC^* distal colon sections (D) and colon whole mounts (E). Scale bars, 100 μm (D) and 200 μm (E). Graphs show quantification of percentage of PAS-positive (PAS^+^) IECs per crypt (D) and percentage of Mucin 2-positive (Muc2^+^) area per whole mount area examined (E). n = 3-4. Data are presented as mean ± SEM, \**P* < 0.05, \*\**P* < 0.01, †*P* < 0.001.

### Plectin-deficient IECs form aberrant cell junctions and disordered KF networks

As the structural and functional integrity of epithelia is secured by cell junctions^2, 29^, we compared the morphology of cell-ECM (HDs) and cell-cell (TJs, AJs, and Ds) adhesions formed by *Ple^fl/fl^* and *Ple^ΔIEC^* IECs, using transmission electron microscopy (TEM). A quantitative analysis of the HD size revealed an extended cross-sectioned length of seemingly less electrodense HD plaques in the *Ple^ΔIEC^* colon; furthermore, the space between HDs and the BM was significantly dilated (Figure 3A). Similar to HDs, we also found significantly dilated intercellular spaces of TJs, AJs, and Ds between adjacent *Ple^ΔIEC^* IECs (Figure 3A). These morphological alterations coincided with generally lower expression levels of the hemidesmosomal constituents Itgα6 and Itgβ4 and the following cell-cell junctional markers: zonula occludens 1 (ZO-1; TJs), E-cadherin (E-cad; AJs), desmoglein 2 (Dsg2; Ds), and desmoplakin 1/2 (Dsp1/2; also Ds) at both mRNA and protein levels (Figure 3B-E). These results clearly show that plectin deficiency leads to the formation of aberrant intestinal junctional complexes, which likely accounts for breached epithelial barrier integrity.

**Figure 3.**
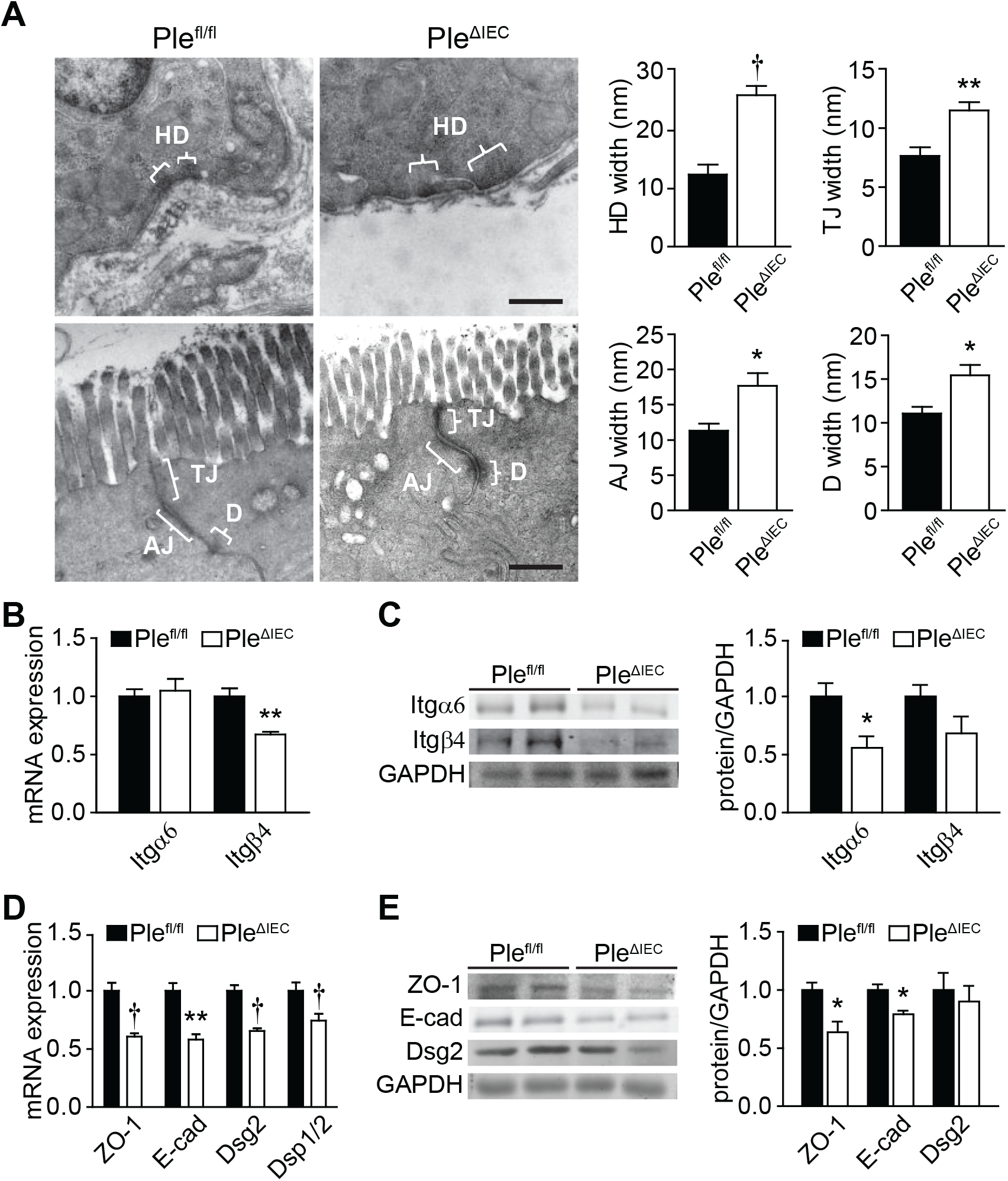
Formation of aberrant cell junctions in *Ple^ΔIEC^* IECs. (A) Representative TEM micrographs of *Ple^fl/fl^* and *Ple^ΔIEC^* IEC junctional complexes. Braces (white) indicate hemidesmosomes (HD), tight junctions (TJ), adherens junctions (AJ), and desmosomes (D). Scale bar, 500 nm. Graphs show quantitative analyses of junctional complex widths (measured as distance from IEC to BM (HD) or distance from IEC to IEC membrane (TJ, AJ, and D)). 5 to 15 junctions were measured (2 mice per genotype). (B) Relative mRNA levels of integrin (Itg) α6 and β4 in scraped mucosa from *Ple^fl/fl^* and *Ple^ΔIEC^* distal colons. n = 4-5. (C) Quantification of Itgβ4 and Itgα6 in scraped distal colon mucosa from *Ple^fl/fl^* and *Ple^ΔIEC^* mice by immunoblotting. GAPDH, loading control. Graph shows relative band intensities normalized to average *Ple^fl/fl^* values. n = 3. (D) Relative mRNA levels of ZO-1, E-cadherin (E-cad), desmoglein 2 (Dsg2), and desmoplakin 1/2 (Dsp1/2) in *Ple^fl/fl^* and *Ple^ΔIEC^* distal colons. n = 5. (E) Quantification of ZO-1, E-cad, and Dsg2 in *Ple^fl/fl^* and *Ple^ΔIEC^* colon mucosa by immunoblotting. GAPDH, loading control. Graph shows relative band intensities normalized to average *Ple^fl/fl^* values. n = 3. Data are presented as mean ± SEM,\**P* < 0.05, \*\**P* < 0.01, †*P* < 0.001.

In previous studies, we showed that plectin controls cell junctions through anchorage of IF networks^20, 22, 30^. Therefore, we next compared the appearance of KFs in *Ple^fl/fl^* and *Ple^ΔIEC^* colon sections using immunofluorescence microscopy. Although the general appearance of K8 and K19 networks did not significantly differ in the two cell types (Figure S5), super-resolution microscopy of pan-K-labelled sections revealed less pronounced apical staining of *Ple^ΔIEC^* IECs (Figure 4A). Moreover, in *Ple^ΔIEC^* IECs, pan-K positive filaments formed less-ordered and rather coarse meshworks, while *Ple^fl/fl^* IECs displayed typical staining patterns with filaments regularly aligned along the apico-basal axis (Figure 4A). The changes in KF network organization were not caused by altered keratin (K8, K18, and K19) expression, as no differences were found at either the mRNA or the protein level (Figure 4B,C). No apparent abnormalities were seen in actin filament and microtubule organization (Figure S5).

**Figure 4.**
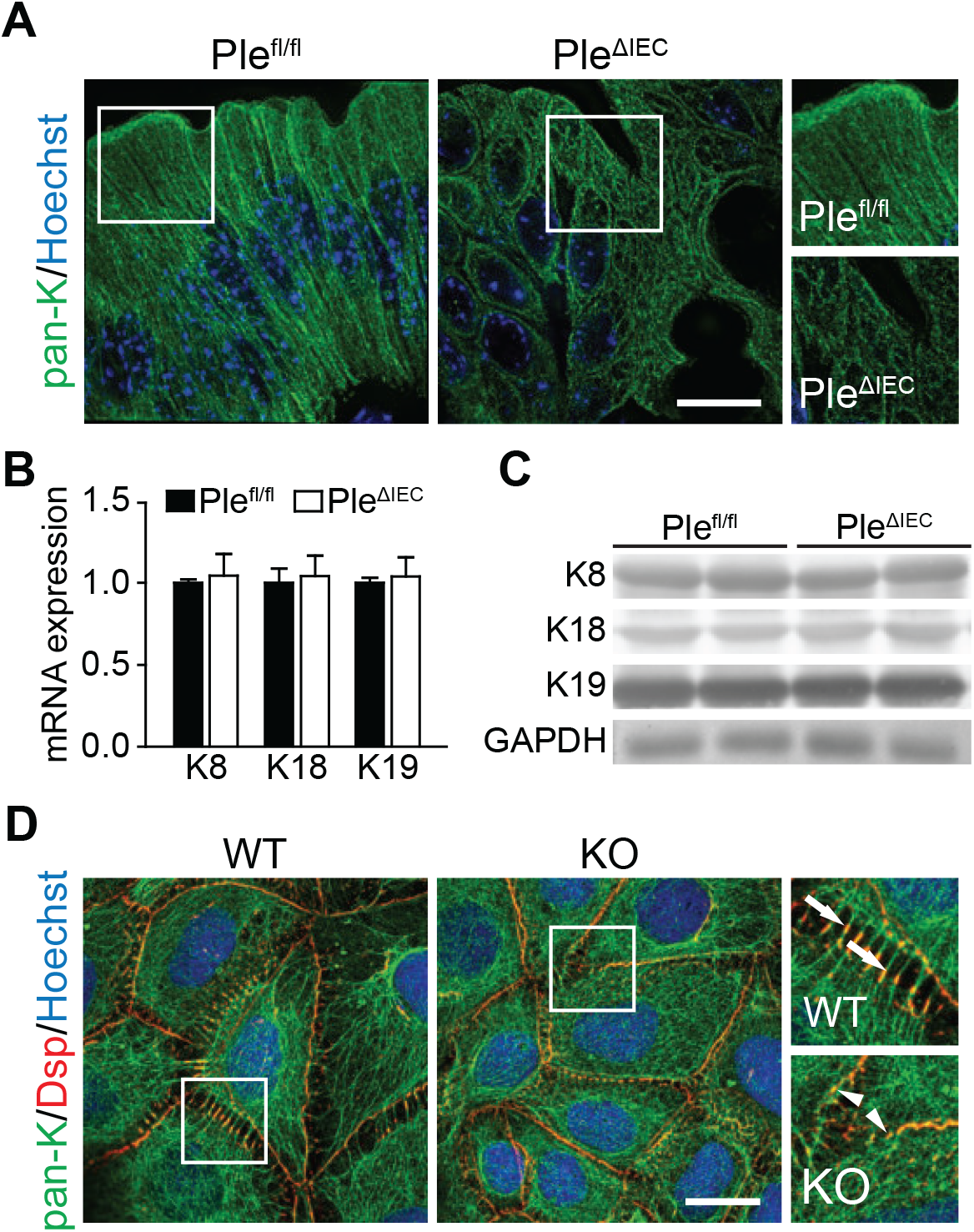
Plectin organizes KFs in IECs. (A) Representative super-resolution STED images of *Ple^fl/fl^* and *Ple^ΔIEC^* distal colon sections immunolabelled for pan-keratin (pan-K; green) with nuclei stained with Hoechst (blue). Scale bar, 10 μm. Boxed areas show ×1.3 images. (B,C) Relative mRNA (B) and protein (C) levels of K8, 18, and 19 in *Ple^fl/fl^* and *Ple^ΔIEC^* distal colon. n = 3-5. Data are presented as mean ± SEM, *P* > 0.05 by unpaired Student *t* test. (D) Representative immunofluorescence images of WT and KO Caco-2 cell monolayer cultures immunolabelled for pan-K (green) and desmoplakin (Dsp; red). Nuclei were stained with Hoechst (blue). Arrows, straight K8 filaments anchored to Dsp-positive desmosomes; arrowheads, tangled K8 filaments. Scale bar, 20 μm. Boxed areas show ×2.5 images.

Aberrant KF cytoarchitecture was also clearly discernible in pan-K immunolabelled monolayers of plectin-deficient (KO) human IECs (Caco-2). To mimick the *in vivo* situation, mature differentiated Caco-2 cells 16 days after the confluency were used. In wild-type (WT) cells, the KF network was densely packed around the cell center, from which individual KFs extended towards the cell periphery delineated by clearly defined desmoplakin-positive Ds (Figure 4D). In contrast, KO cells showed tangled KFs, which were evenly distributed throughout the cytoplasm and seemingly overlapped with rather continuous desmoplakin-positive structures at the cell-cell borders (Figure 4D). Collectively, these findings indicate that plectin ablation in IECs results in altered keratin network organization and aberrant KF anchorage to desmosomal junctions.

### Plectin preserves intestinal epithelial integrity through HD stabilization

Plectin-mediated attachment of the keratin network to Itgα6β4-containing HDs plays a crucial role in stabilizing the adhesion of keratinocytes to the matrix and hence imparts mechanical stability to the skin^20, 31^. To examine whether the *Ple^ΔIEC^* intestine phenotypically follows the same paradigm, we scrutinized colon and small intestine sections immunolabelled for K8 and Itgα6 (Figure 5A) or collagen (Col) IV (Figure S6). In line with the observations from H&E- and Sirius red-stained colon sections (Figures 2A and S6), *Ple^ΔIEC^* IECs partially lost their polarized orientation; they were misaligned and largely detached from the BM at the luminal surface of the crypts (Figure 5A, upper panels). Extensive detachment of *Ple^ΔIEC^* IECs was even more apparent in the small intestine, where we often found the whole epithelial sheet physically separated from underlying structures (Figure 5A, lower panels). Remarkably, in both the *Ple^ΔIEC^* colon and the small intestine, Itgα6-positive patches remained confined to the BM, while detached IECs were entirely devoid of Itgα6 signals. Thus we conclude that plectin ablation abrogates the functional link between KFs and HDs.

**Figure 5.**
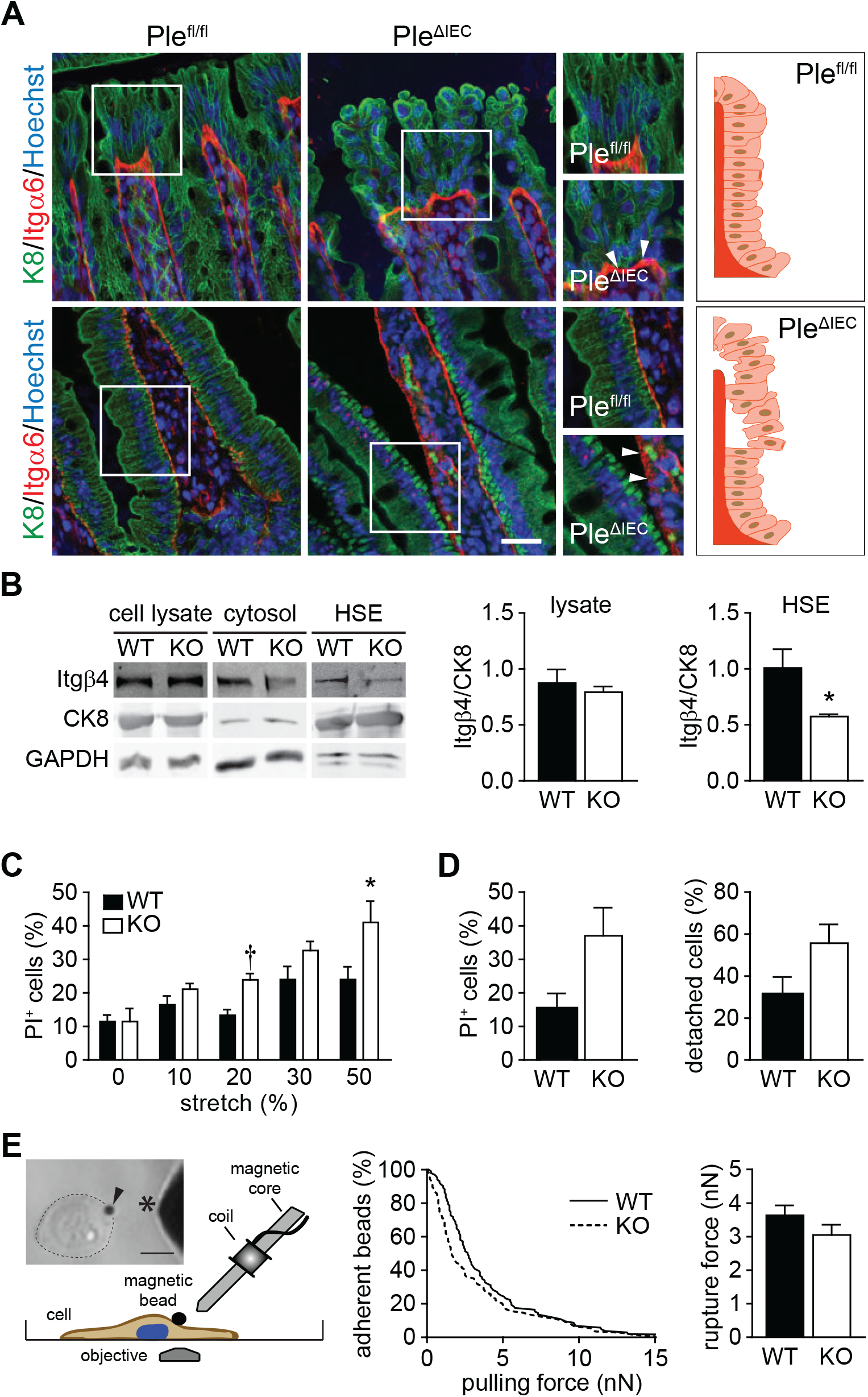
Plectin stabilizes IEC hemidesmosomes through KF recruitment. (A) Representative immunofluorescence images of *Ple^fl/fl^* and *Ple^ΔIEC^* distal colon (upper panels) and small intestine (lower panels) sections immunolabelled for K8 (green) and Itgα6 (red); Hoechst-stained nuclei (blue). Arrowheads, Itgα6-positive clusters. Scale bar, 25 μm. Boxed areas show ×1.5 images. Drawn schemes depict aligned, BM-attached *Ple^fl/fl^* IECs (upper panel) and mislocalized, detaching *Ple^ΔIEC^* IECs (lower panel). (B) Cell lysates, cytosol fractions, and keratin-enriched high salt extracts (HSE) were prepared from WT and KO Caco-2 cells and subjected to immunoblotting with antibodies to Itgβ4 and CK8. GAPDH, loading control. Graphs show relative band intensities normalized to average *Ple^fl/fl^* values. n = 4-6. (C) Viability of WT and KO Caco-2 cells exposed to uniaxial cyclic stretch presented as percentage of dead (PI-positive; PI^+^) cells. n = 9-11. (D) Quantification of WT and KO Caco-2 cell viability (left) and adhesion (right) under radial shear flow shown as percentage of dead and detached cells, respectively. n = 6. (E) Adhesion strength between ECM-coated paramagnetic beads and WT and KO Caco-2 cells was quantified using magnetic tweezers. Image and schema depict magnetic tweezer setup. Arrowhead, paramagnetic bead; asterisk, magnetic tweezer tip; dotted circular line, cell border. Scale bar, 20 μm. Curve graph shows quantification of bead detachment (given as percentage of adherent beads at given pulling force), and bar graph shows cumulative rupture force (calculated from median bead detachment. n = 103. Bar graph data represent mean ± SEM, \**P* < 0.05, †*P* < 0.001.

To address the effects of plectin ablation biochemically, we prepared keratin-enriched cell fractions^19^ from human WT and KO IEC Caco-2 lines and compared their integrin content by immunoblotting using antibodies to Itgβ4. As expected, such cell fractions were highly enriched in keratins 8 (Figure 5B), 18, and 19 (not shown). Although Itgβ4 levels were comparable in cell lysates, the Itgβ4 content of insoluble keratin fractions was significantly reduced in KO cells compared to WT cells (Figure 5B). These observations correlated well with the histological data (Figure 5A).

To assess whether plectin deficiency affects biomechanical properties of IECs, we performed a series of quantitative assays with human IEC lines Caco-2 and hCC. Monitoring cell viability under mechanical stress on a stretched flexible membrane (uniaxial cyclic stretch) revealed higher mechanical vulnerability of both KO cell lines, as the proportion of PI-positive (dead) cells significantly increased with stretch amplitudes ranging from 10% to 50% (Figure 5C and Figure S7A). Significantly reduced mechanical resistance of KO cells was confirmed by fluid shear stress assay using a spinning disc device. When exposed to constant radial flow, KO cells displayed death rates about twice as high as that of their WT counterparts (Figure 5D and Figure S7B). Moreover, the fraction of detached Caco-2 (but not hCC) cells was higher for KO than WT cells. This suggests that plectin ablation weakens their adhesion to the underlying substratum.

To confirm this hypothesis, we quantified adhesion strength between ECM-coated superparamagnetic beads and cell adhesions using magnetic tweezers. We then applied increasing forces of up to 15 nN and recorded the force at which each bead detached (ruptured) from the cell. From a total of >100 rupture events for each cell type, we calculated the cumulative detachment probability as a function of pulling force and report the rupture force at which 50% of the beads detached from the cells (Figure 5E and Figure S7C). As expected, we measured lower forces in both Caco-2 and hCC KO compared to WT cells, which confirms our hypothesis of weaker Itg-mediated adhesions in plectin-deficient cells. Hence, like for skin type I HDs ^20^, plectin loss is deleterious for the stability of type II HDs present in the intestine, leading to compromised mechanical resilience of IECs and intestinal epithelia.

### IEC-specific plectin deficiency exacerbates experimental colitis

The spontaneous colitic phenotype in *Ple^ΔIEC^* mice (Figure 1) suggests that plectin deletion can contribute substantially to the pathogenesis of UC. To assess whether loss of plectin increases the susceptibility to colitis, we induced experimental colitis in *Ple^fl/fl^* and *Ple^ΔIEC^* mice. Even a short exposure (3-4 days) to low DSS doses (1.5-2%) resulted in a dramatic body weight loss of *Ple^ΔIEC^* mice, in sharp contrast to similarly treated *Ple^fl/fl^* which experienced only insignificant weight losses (Figure 6A). The weight loss of mutant mice was associated with a higher disease activity index (DAI; Figure 6A), a decreased survival rate (not shown), and a significant reduction in colon length (Figure 6B). The severity of induced colitis in *Ple^ΔIEC^* mice coincided with larger inflamed areas at days 4 and 6 after the initiation of DSS-treatment (Figure 6C), corresponding to a higher influx of MPO-positive neutrophils and intestinal epithelial injury. Further, histological evaluation of ‘Swiss rolls’ of the entire colon confirmed these results and revealed clear signs of inflammation and epithelial damage in all DSS-treated animals. However, large regions with heavy ulceration, crypt damage, and inflammatory response in *Ple^ΔIEC^* mice were in striking contrast to fewer lesions in *Ple^fl/fl^* mice (Figure 6D and Figure S8A).

**Figure 6.**
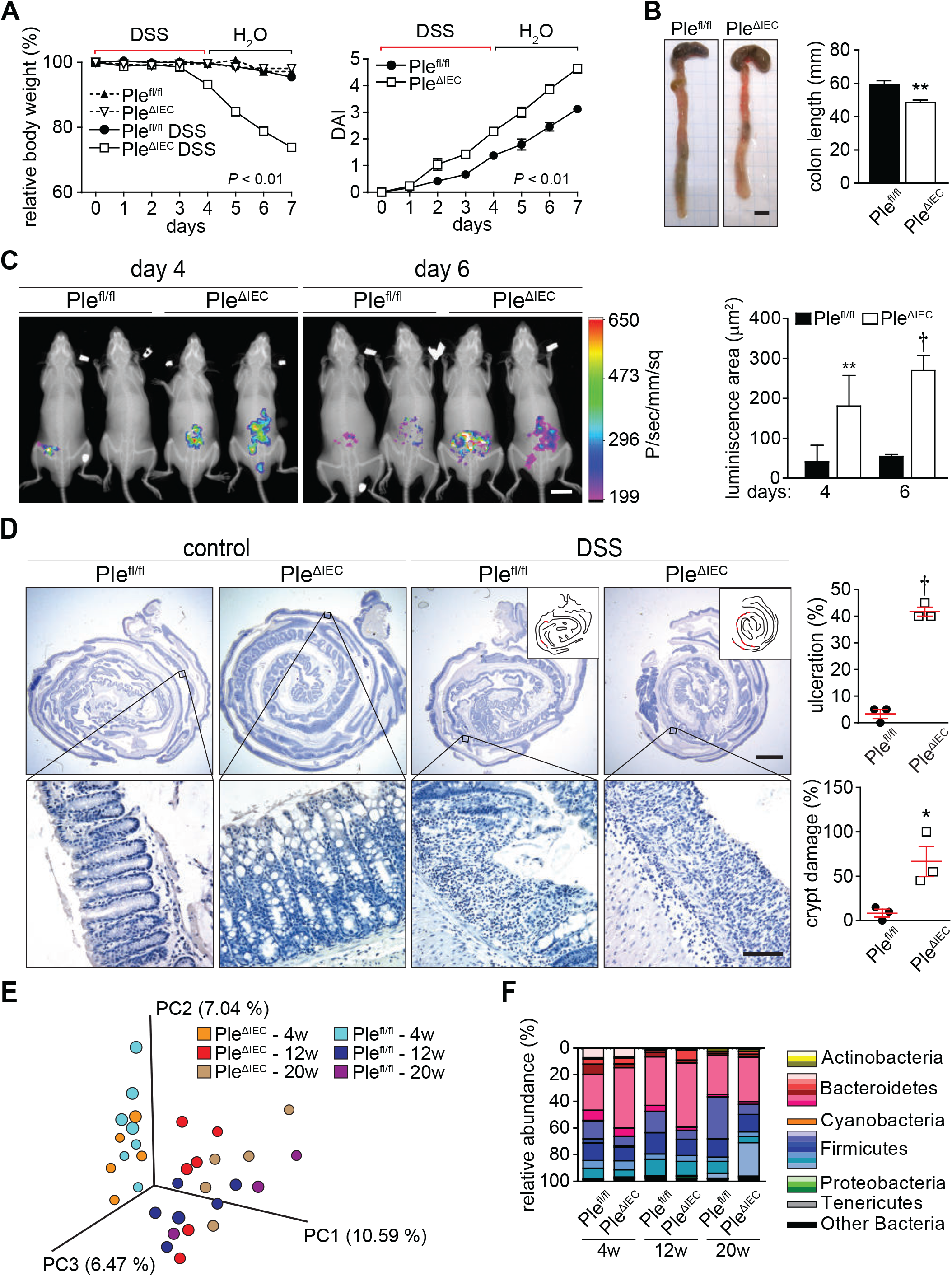
*Ple^ΔIEC^* mice are more susceptible to DSS-induced colitis. (A) Relative body weight and disease activity index (DAI) of untreated and DSS-treated *Ple^fl/fl^* and *Ple^ΔIEC^* mice during experimental colitis. 4-7 mice per genotype and time point were analyzed. (B) Representative images of colon and caecum of DSS-treated *Ple^fl/fl^* and *Ple^ΔIEC^* mice. Graph shows colon length. n = 4-6. (C) *In vivo* chemiluminescence images and signal quantification (graph) of DSS-treated *Ple^fl/fl^* and *Ple^ΔIEC^* mice injected with the myeloperoxidase substrate luminol on day 4 and 6 of DSS treatment. n = 3-4. (D) Representative hematoxylin-stained sections of Swiss roll mounts from untreated (control) and DSS-treated (DSS) mice. Scale bars, 2 mm, magnified boxed areas, 100 μm. Insets, outlines of lesions (in red) distributed along mucosa (black lines) in corresponding panels. Graphs show quantification of colonic tissue damage given as percentage of ulceration and crypt damage. n = 3. (E, F) Fecal microbiota beta diversity in 4-, 12-, and 20-week-old untreated *Ple^fl/fl^* and *Ple^ΔIEC^* mice as determined by 16S rDNA sequencing. Principal Coordinate Analysis plot (E), constructed with unweighted UniFrac distance metric, shows clustering of microbial beta diversity. PC1, PC2, and PC3 represent top three principal coordinates that captured most of diversity (given as percentage). Global composition (F) of bacterial microbiota at phyla level shown as relative operational taxonomic unit (OTUs) abundance per time point and genotype. n = 4-6. Data are presented as mean ± SEM, \**P* < 0.05, \*\**P* < 0.01, †*P* < 0.001.

Since gut microbial dysbiosis is a typical finding in UC patients^32–34^, we compared the composition of fecal microbiota in unchallenged *Ple^fl/fl^* and *Ple^ΔIEC^* mice at the ages of 4, 12, and 20 weeks. Surprisingly, despite the impaired intestinal barrier and concomitant inflammation phenotype of *Ple^ΔIEC^* mice (Figure 1F-L), we observed no significant differences in alpha (Figure S8B) and beta (Figure 6E) diversities between both genotypes. In all animals, bacterial microbiota were dominated by bacteria belonging to families S24-7 (Bacteroidetes), Lactobacillaceae (Firmicutes), and Lachnospiraceae (Firmicutes) (Figure 6F). Together, these data show higher susceptibility of *Ple^ΔIEC^* mice to DSS-induced colitis, accompanied by severe epithelial damage and inflammation in the absence of microbial dysbiosis.

### Reduced mechanical stability of epithelia accounts for intestinal injury in *Ple^ΔIEC^* mice

To identify the onset and time course of intestinal injury in *Ple^ΔIEC^* mice, we assessed intestinal epithelial damage scores in newborn, 21-day-old, and 12-week-old mice (Figure S9A and Figure 7A-C). While newborn mice were histologically inconspicuous, the colon and the small intestine displayed first signs of damage in 21-day-old *Ple^ΔIEC^* mice, which coincided with weaning and transition to solid chow. To gain better control over *plectin* inactivation timing, we generated TMX-inducible IEC-specific *plectin* knockout (*Ple^ΔIEC-ERT2^*) mice. Three consecutive applications of TMX in 9-week-old *Ple^ΔIEC-ERT2^* mice resulted in recombination efficiency comparable to that of constitutive *Ple^ΔIEC^* mice, and a distinctive intestinal phenotype developed as early as 5 days post-TMX administration (not shown). To determine the effect of a diet change on intestinal injury, *Ple^ΔIEC-ERT2^* and *Ple^fl/fl^* control mice were either kept on solid chow or provided with a low-residue liquid diet 6 days before TMX administration. The liquid diet significantly attenuated epithelial damage in the colon of *Ple^ΔIEC-ERT2^* mice; however, the histological score indicates more severe injury of the small intestine (Figure 7D-F and Figure S9B). The beneficial effects of the liquid diet also manifested as less prominent colon swelling (not shown). As previous studies linked the severity of colitis and intestinal injury with commensal microbiota^8, 35^, next we treated *Ple^fl/fl^* and *Ple^ΔIEC-ERT2^* mice with well-established broad-spectrum antibiotics^8^. In our experimental setup, the apparent milder colitic phenotype consistently coincided with lower epithelial damage in colon of *Ple^ΔIEC-ERT2^* mice. This treatment however did not affect epithelial injury of the *Ple^ΔIEC-ERT2^* small intestine (Figure 7G-I and Figure S9C). Collectively, these data support the notion that the increased susceptibility of the plectin-deficient intestinal epithelium to mechanical strain impinged by luminal content is caused by a lack of KF attachment to integrin clusters and destabilization of intestinal HDs. Further, the fact that antibiotics also partially alleviate epithelial damage suggests that luminal bacteria significantly contribute to intestinal injury in *Ple^ΔIEC-ERT2^* _mice._

**Figure 7.**
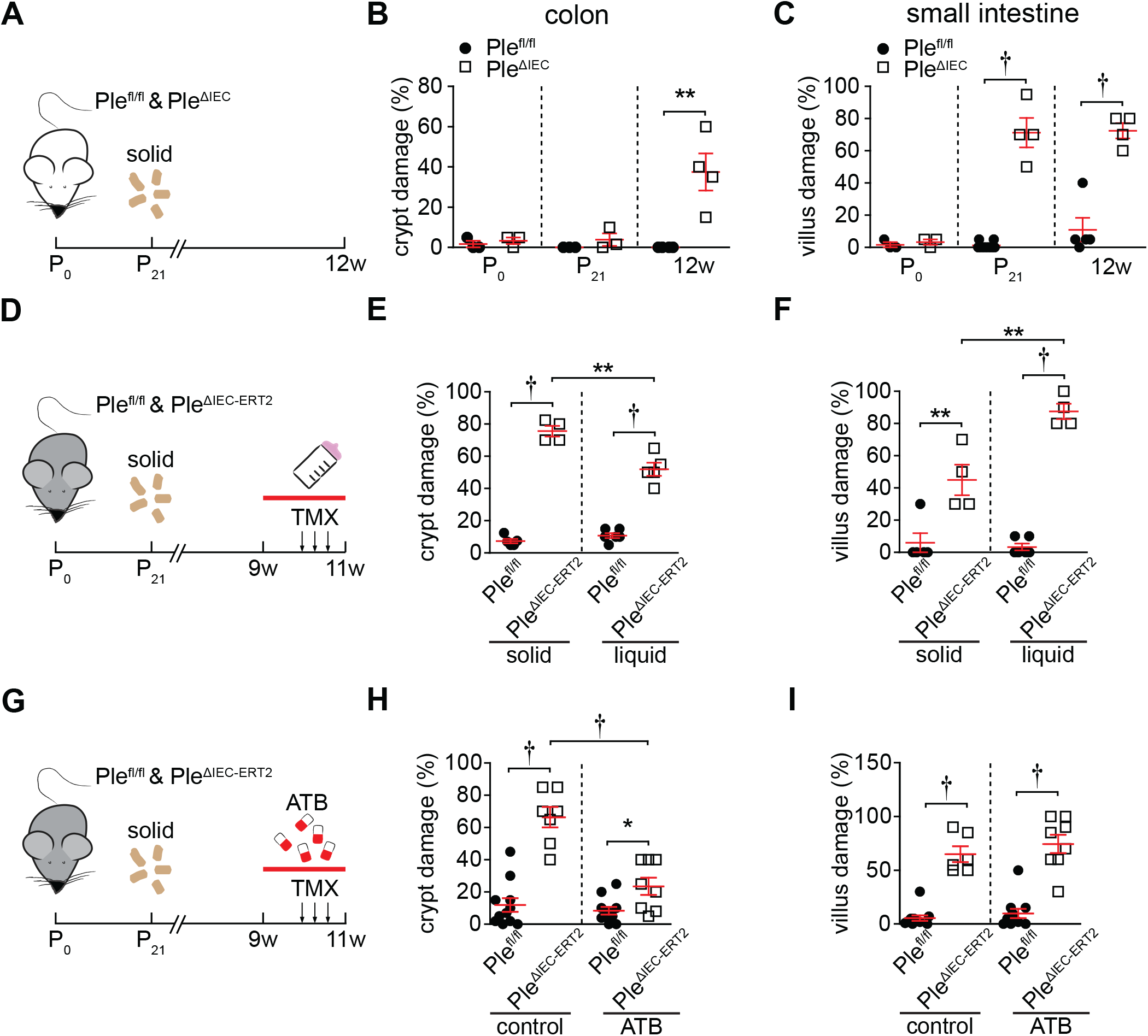
Intestinal epithelial damage in *Ple^ΔIEC^* mice results from mechanical stress. (A-C) *Ple^fl/fl^* and *Ple^ΔIEC^* mice were sacrificed on postnatal day 0 (P0), postnatal day 21 (P21), and at 12 weeks (12w) of age, and epithelial damage scores were assessed from colon and small intestine sections. Schematic illustrates experimental setup (A). Solid, transition to solid chow at P21. Graphs show quantification of epithelial damage in colon (B) and small intestine (C) at age indicated. (D-F) 9-week-old *Ple^fl/fl^* and *Ple^ΔIEC-ERT2^* mice were either kept on solid chow or provided with liquid diet for 14 days. *Plectin* inactivation was induced by three consecutive applications of tamoxifen (TMX) on days 6, 8, and 10; mice were sacrificed on day 14. Schematic illustrates experimental setup (D). Solid, transition to solid chow at P21; arrows, TMX application; red bar, period on liquid diet. Graphs show quantification of epithelial damage in colon (E) and small intestine (F) on solid chow and liquid diet. (G-I) 9-week-old *Ple^fl/fl^* and *Ple^ΔIEC-ERT2^* mice were kept either untreated or treated with broad-spectrum antibiotics. *Plectin* inactivation and sample collection were identical to B. Schematics ilustrate experimental setup (G). Chow, transition to solid chow at P21; arrows, TMX application; red bar, period of antibiotics (ATB) treatment. Graphs show quantification of epithelial damage in colon (H) and small intestine (I) on solid chow and liquid diet. Data are presented as mean ± SEM, \**P* < 0.05, \*\**P* < 0.01, †*P* < 0.001.

## DISCUSSION

The intestinal epithelium faces substantial mechanical stress^36^, which is inextricably linked to gut physiology. Although several studies suggest the importance of intestinal KF networks^35, 37, 38^ and KF-associated cell junctions (Ds and HDs)^8, 9^ for protection against intestinal inflammation and CRC, the contribution of altered epithelial mechanics to observed phenotypes remain unexplored. Here, we focus on the role of KF-cell junction linker plectin in the maintenance of intestinal homeostasis, and we provide a comprehensive analysis of molecular mechanisms governing the mechanical stability of intestinal epithelia.

The most notable phenotype of both plectin-deficient mouse models (*Ple^ΔIEC^* and *Ple^ΔIEC-ERT2^*) is the detachment of IECs from the underlying BM, resulting in extensive epithelial injury and eventually in the spontaneous development of a colitic phenotype. Strikingly, this is accompanied by loss of the hemidesmosomal ECM receptor Itgα6 from detached IECs, while Itgα6 patches remain on a collagen-stained BM, likely indicating their inefficient linkage to cytoskeletal structures. Indeed, our TEM analysis revealed that less electrodense HDs formed by *Ple^ΔIEC^* IECs were somewhat elongated, and gaps between HD plaques and the BM were significantly wider compared to *Ple^fl/fl^* IECs. The formation of morphologically abnormal HDs was paralleled with reduced expression levels of both HD-forming integrins (α6 and β4) in the *Ple^ΔIEC^* mucosa. Moreover, the content of Itgβ4 was also significantly diminished in keratin-enriched fractions prepared from plectin-deficient IECs, suggesting reduced association of Itgα6β4 complexes with intestinal keratins (K8, 18, and 19).

Our observations are concordant with a recently published model for skin type I HD^10^, which proposed that plectin (along with BPAG, another plakin family member) fortifies HD plaques both horizontally (by a lateral association of Itgβ4) and vertically (by interlinking Itgβ4 with KFs). Accordingly, ablation of plectin, the only plakin present in intestinal type II HD^39^, would fully abrogate a functional link between KFs and HDs and would result in their overall destabilization. In line with this hypothesis, we show *in vitro* higher detachment (paralleled with a higher death rate) of plectin-deficient IECs exposed to a uniaxial cyclic stretch and a constant radial flow compared to their WT counterparts. Consistently, we also determine considerably lower adhesion strength between ECM-coated superparamagnetic beads and plectin-deficient IECs using magnetic tweezers. Hence, by combining *in vivo* and *in vitro* approaches, we provide evidence that plectin is essential for the stability of intestinal HD type II, a structure preventing colitis ^8^ and presumably also the risk of colitis-associated CRC^8, 40, 41^.

Previous studies demonstrated that the deletion of *plectin* has adverse effects on the formation of intercellular junctions, with consequences for the epithelial barrier function^21, 22^. It has been shown that plectin-deficient cholangiocytes form dysfunctional Ds and fail to upregulate some desmosomal proteins, such as desmoplakin, a putative binding partner of plectin^42^, in response to bile stasis^22^. This failure results in mechanical weakening of the biliary epithelium and contributes to plectin-related familial intrahepatic cholestasis^43^. In terms of mechanistic parallels between plectin-deficient biliary and intestinal epithelia, we found that apart from destabilizing HDs, plectin deficiency also leads to prominent broadening of Ds, AJs, and TJs. Moreover, *Ple^ΔIEC^* IECs exhibit downregulation of corresponding junctional constituents (ZO-1, E-cad, Dsg2, and Dsp1/2). Although the resulting dilatation of intercellular spaces would *per se* suffice to explain the observed increase in intestinal permeability and bacterial penetration, the “leaky gut” in *Ple^ΔIEC^* mice seems ultimately rooted in the less firm IEC/BM connection, given the extent of IEC detachment. On the other hand, proper anchorage of KFs (determining cell mechanics) to Ds (ensuring intercellular cohesion) is known to provide load-bearing tissues with mechanical stability ^11^. Showing altered KF cytoarchitecture and aberrant D formation in both *in vitro* plectin-deficient IEC systems and *Ple^ΔIEC^* mice, our results suggest that D-keratin complex abnormality substantially contributes to the compromised mechanics of the *Ple^ΔIEC^* intestinal epithelium.

We propose that the lack of functional plectin at HDs (in combination with its effects on KFs and Ds) and the resulting mechanical epithelial fragility favor an impaired intestinal barrier function and are ultimately responsible for colitis. Importantly, comparable mucosal deterioration was observed upon plectin ablation during development and in the adult intestine with its fully mature immune system. Furthermore, we also demonstrate that loss of plectin can exacerbate experimental colitis in mice. Consistent with these results, lower expression levels of plectin correlate with UC development in human patients, suggesting that defects in cytoskeleton coordination mediated through plectin contribute to IBD pathogenesis in humans by affecting IEC/BM adhesion, IEC cohesion, and mechanical properties. However, it is well recognized that properly organized KF networks^38, 44^, HDs^8^, and intercellular junctions^2, 3, 9^ exert numerous non-mechanical functions, providing the intestinal epithelium with protection against microbial infection and uncontrolled inflammation. Our data do not rule out similar functions in the *Ple^ΔIEC^* intestine. Further studies will be required to investigate how plectin deficiency affects cell-autonomous (barrier function-independent) mechanisms involved in the interplay between IECs, gut microbiota, and immune cells.

The rapid deterioration of the *Ple^ΔIEC^* intestinal mucosa following weaning (i.e., a switch to a solid diet and the amplification of muscle contractions) indicates that the origin of colitis in the absence of plectin is primarily associated with a reduced capacity of IECs to resist mechanical stress. In addition, the most severe epithelial injury was found in the distal colon, which is the region most intensely subjected to such stress. Comparably devastating epithelial instability has been well documented for epidermal layers in EB patients^15, 16^. As there is no causal therapy for EB available^15^, the current treatment focuses mainly on prevention of tissue destruction. Following the same rationale, we demonstrate that a low-residue liquid diet significantly attenuates colonic epithelial damage, thus protecting its barrier function. Surprisingly, this approach aggravates IEC detachment in the *Ple^ΔIEC^* small intestine, which might suggest augmented susceptibility of the small intestine to plectin loss. Therefore, future studies should investigate whether differential expression of plectin along the gastrointestinal tract might have an impact on regional differences in disease manifestations in patients. In line with previous observations^8, 35^, antibiotic treatments markedly decrease not only mucosal inflammation but, intriguingly, also epithelial damage, which implies that host-microbiota interactions contribute to excessive IEC sloughing in the *Ple^ΔIEC^* intestine. Although dietary factors can likely ameliorate only less extensive trauma, our results suggest that a low-residue liquid diet combined with antibiotic treatment might be a useful palliative modality. To translate our findings into clinical medicine: it remains to be determined whether such a strategy i) is suitable for long-term treatment and ii) is effective with respect to systemic disease manifestation.

## METHODS

### Supplemental methods are available online

#### Patients

Colon biopsy samples were collected from patients with active UC (n = 51) and from healthy controls (n = 18) admitted to the Hepatogastroenterology Department at the Institute for Clinical and Experimental Medicine (Prague, Czech Republic) for a colonoscopy from July 2016 to May 2019. Subjects were assigned to the healthy control group only after all clinical examinations excluded any signs of autoimmune disease, inflammatory disease, and colon cancer. All UC patients with concurrent primary sclerosing cholangitis (PSC) were excluded from the study. Endoscopic UC activity at the time of a standard optical colonoscopy was categorized according to the Mayo endoscopic subscore and confirmed by histology examinations of the grade of inflammation. Clinical characteristics of patients are shown in Supplementary Table S1. Standard endoscopic biopsies were extracted from the inflamed non-dysplastic mucosa of the left colon (rectum) and immediately placed in an RNAlater solution. Total RNA was extracted according to the manufacturer’s instructions.

#### Mice

*Plectin^flox/flox^* (*Ple^fl/fl^*) mice^23^ were crossed with *villin-Cre* transgenic mice (MGI 2448639) to generate *Ple^fl/fl^*/*villin-Cre* mice (*Ple^ΔIEC^*) and with *villin-creERT2* transgenic mice (MGI 3053826; both Cre strains were kindly provided by S. Robine^45^) to generate *Ple^fl/fl^*/*villin-creERT2* mice (*Ple^ΔIEC-ERT2^*). Age-matched littermate male mice were used in all experiments. Unless stated otherwise, mice were 12-14 weeks old. Animals were housed under specific pathogen-free conditions with regular access to chow and drinking water and a 12 hours light/12 hours dark regime.

#### Cells and CRISPR-mediated targeting of plectin

Caco-2 cells were grown in Dulbecco’s modified Eagle medium (DMEM) supplemented with 20% fetal bovine serum (FBS) in 5% CO_2_/air humidified atmosphere at 37°C. Human colonic cells (hCC; T0570, Applied biological materials, Inc.) were cultured in DMEM supplemented with 10% FBS in 5% CO_2_/air humidified atmosphere at 37°C. Plectin knockout (KO) cell lines were generated by targeting genomic sequences of intron 25-26 and exon 31 of *Plectin* using CRISPR/Cas9 plasmid pX330 Cas9-Venus (a kind gift of B. Schuster, IMG CAS, Prague, Czech Republic) as described previously^22^. The potential off-target sites were predicted using CRISPOR (http://crispor.tefor.net/). The 4 top-ranking potential off-target sites for each guide RNA were selected for validation. The genomic DNA sequences surrounding the potential off-target sites were amplified by PCR using gene-specific primers (Supplementary Table 3). PCR products were analyzed by direct sequencing (Figures S12-13).

#### Statistics

All results are presented as mean ± SEM. All normally distributed parametric data were analyzed by 2-tailed unpaired Student *t* test. Comparisons of multiple groups to controls were performed using 2-tailed 1-way ANOVA. Comparisons of frequency distributions of BrdU-positive cells were analyzed with Mann-Whitney test. Survival curves were analyzed by Mantel-Cox test. Statistical analyses were performed using GraphPad Prism 5 (GraphPad Software, Inc., La Jolla, CA). Statistical significance was determined at the levels of \**P* < 0.05, ** *P* < 0.01, † *P* < 0.001; n values are specified in the figure legends.

#### Study approval

This study was approved by the Ethics Committee of the Institute for Clinical and Experimental Medicine and Thomayer Hospital with Multi-Center Competence (G16-06-25). Written informed consents were obtained from all subjects before the study. All animal studies were performed in accordance with European Directive 2010/63/EU and were approved by the Czech Central Commission for Animal Welfare (48/2014).

## Author contribution

Study concept and design: M.G. Acquisition of data: A.K., P.B., L.S., G.O.-E., G.K., J.Pr., M.S., Ch.D., Z.S., M.V., N.Z. Analysis and interpretation of data: A.K., M.G., J.S., E.S., L.B., J.Pa., J.Pr., D.J., M.K., B.F. Drafting of the manuscript: A.K., M.G. Critical revision of the manuscript for important intellectual content: all authors. Funding: M.G., M.K., G.W., B.F. Technical and material support: P.W., L.B., V.K., R.S., G.W.

## Acknowledgements

We would like to thank L. Macurek (IMG CAS, Prague) for generously providing the hCC cell line, S. Robine (Institut Curie, Paris) for Cre mouse lines, and B.Schuster (IMG CAS, Prague) for pX330 Cas9-Venus plasmid; H. Havelkova (IMG CAS, Prague) and I. Muricova (Institute of Physiology CAS, Prague) for their outstanding technical assistance; S. Reipert (University of Vienna, Vienna), J. Tureckova (IMG CAS, Prague), and Z. Jiraskova (Institute of Microbiology CAS, Prague) for their expertise; and T. Epp, M. Prechova (both IMG CAS, Prague) and A.I. Ivanov (Lerner Research Institute of Cleveland Clinic Foundation, Cleveland) for critical reading of the manuscript. We acknowledge the Light and Electron Microscopy Core Facilities, IMG CAS, Prague, Czech Republic, for their support with advanced microscopy imaging.

## SUPPLEMENTARY MATERIAL

### 1. Supplementary METHODS

#### DSS-induced colitis and disease activity scoring

12-week-old *Ple^fl/fl^* and *Ple^ΔIEC^* mice were provided with a 2% DSS (TdB Consultancy) dissolved in drinking water *ad libitum* over 4 days. Then they were provided with drinking water over 3 days. Mice were sacrificed on day 8. As described previously^1^, body weight, stool consistency, and rectal bleeding (Hemoccult Fecal Occult Blood Test, Beckman Coulter) were assessed daily to calculate the disease activity index (DAI).

#### Depletion of gut microbiota by antibiotic treatment

Streptomycin 2 g/l (Carl Roth), gentamycin 200 mg/l (Carl Roth), enrofloxacin 100 mg/l (Sigma-Aldrich), and bacitracin 1 g/l (Carl Roth) were provided to 9-week-old *Ple^fl/fl^* and *Ple^ΔIEC ERT^*^2^ mice in drinking water *ad libitum* over 2 weeks. To induce specific plectin deletion in 9-week-old *Ple^ΔIEC ERT^*^2^ mice, 5 mg of tamoxifen (Sigma-Aldrich; dissolved in 200 μl of sunflower oil) was administered twice a day on days 6, 8, and 10 of the antibiotic treatment by orogastric gavage. The stool was collected before and after the treatment, and its microbial content was analyzed. Mice were sacrificed on day 14 of the treatment. *Ple^fl/fl^* mice received sunflower oil only.

#### Liquid diet feeding

9-week-old *Ple^fl/fl^* and *Ple^ΔIEC ERT^*^2^ mice received a low-residue Ensure Plus (Abbott Laboratories) nutritional supplement diluted 1:1 in drinking water in the course of 2 weeks *ad libitum* in the absence of solid chow. Water was offered *ad libitum*. 5 mg of tamoxifen (Sigma-Aldrich; dissolved in 200 μl of sunflower oil) was administered twice a day on days 6, 8, and 10 of the liquid diet by orogastric gavage. Mice were sacrificed on day 14 of the liquid diet. Control mice received solid chow over the same time period. *Ple^fl/fl^* mice received sunflower oil only.

#### BrdU incorporation assay

14-week-old *Ple^fl/fl^* and *Ple^ΔIEC^* mice received an intraperitoneal injection of 50 mg/kg 5-bromo-2’-deoxyuridine (BrdU; Sigma-Aldrich). Mice were sacrificed 2, 24, and 48 hours after the injection. The small intestine and the colon were dissected and processed for immunohistochemistry (see below). BrdU-positive cells were visualized by anti-BrdU antibody (BMC9318, Roche).

#### Whole-body imaging of inflammation

Myeloperoxidase (MPO) activity was detected by *in vivo* whole-body imaging using a specific XenoLight RediJect Chemiluminescent Inflammation Probe (PerkinElmer). A freshly thawed probe solution was administered by an intraperitoneal injection (200 mg/kg). Mice were immediately moved into the imaging chamber (Xtreme – whole-body imaging system, Bruker), and the number of photons produced by an inflammation probe was recorded (10 min after its administration) in anesthetized mice with 5 min exposure time.

#### Histology, immunohistochemistry, and immunofluorescence

Formalin-fixed and paraffin-embedded small intestine (ileum) and colon sections (5 µm) were stained with haematoxylin and eosin (H&E; Sigma-Aldrich), Sirius Red^2^, and the periodic acid-Schiff stain (Sigma-Aldrich) according to the manufacturer’s protocol. For immunohistochemistry and immunofluorescence, paraffin sections were subjected to heat-induced antigen retrieval in either Tris-EDTA (pH 9) or citrate (pH 6) buffer supplemented with Tween 20 and further permeabilized with 0.1 M glycine and 0.1% Triton X-100 for 15 min. To block endogenous peroxidase activity and non-specific antigen interactions, sections were incubated with 0.3% hydrogen peroxide for 15 min followed by 5% bovine serum albumin (BSA; Sigma-Aldrich) in PBS supplemented with 0.1% Tween 20 (PBST) for 1 hour. Afterwards, sections were incubated with primary antibodies at 4°C overnight, followed by incubation with horseradish peroxidase- (HRP) or fluorophore-conjugated secondary antibodies at room temperature for 1 hour. The HRP signal was visualized with 3,3’-diaminobenzidine (DAB) detection kit (Roche). Caco-2 cells seeded on coverslips were fixed with ice-cold methanol for 1 min. Non-specific antigen interactions were blocked with 5% BSA in PBST for 1 hour. Incubation with primary antibodies followed by incubation with secondary antibodies was performed at room temperature for 1 hour. The following primary antibodies were used: Ki-67 (GTX16667, GeneTex), keratin 8 (Troma I, Developmental Studies Hybridoma Bank), collagen IV (2150-1470, BioRad), pan-keratin (z0622, Agilent/Dako), E-cadherin (610181, BD Biosciences), desmoglein (611002, Progen), desmoplakin (651109, Progen), integrin α6 (ab181551, Abcam), plectin (GP21, Progen), chromogranin A (ab15160, Progen), keratin 20 (clone Ks20.8, Agilent/Dako), keratin 19 (Troma III, Developmental Studies Hybridoma Bank), and β-actin (A2066, Sigma-Aldrich). The following secondary antibodies were used: donkey anti-guinea pig Alexa Fluor (AF) 488, donkey anti-guinea pig AF594, donkey anti-rabbit AF488, goat anti-rabbit AF594, goat anti-rabbit HRP-conjugated, donkey anti-mouse AF488, donkey anti-mouse RhodamineRedX, donkey anti-rat AlexaFluor488, goat anti-mouse HRP-conjugated (all from Jackson ImmunoResearch). Nuclei were counterstained with Hoechst 33258 (Sigma-Aldrich). HRP-conjugated antibodies were visualized by DAB using the DAB Substrate (Roche Diagnostics) and counterstained with Mayer’s hematoxylin (Sigma-Aldrich).

#### Visualization of Muc2 in colonic whole mounts

Staining was performed as described before^1^. Briefly, the intact colon was immediately fixed in Carnoy’s fixative, and cut-open samples (5 x 8 mm) were incubated with an anti-Muc2 antibody (sc-15334, Abcam), at 4°C overnight followed by 1.5 hour-incubation with fluorescently labelled secondary antibodies (donkey anti-rabbit AF488, Jackson ImmunoResearch) at room temperature. Nuclei were counterstained with Hoechst 33258 (Sigma-Aldrich). Samples were placed into 100% glycerol and immediately visualized using a Leica TCS SP8 confocal microscope.

#### TUNEL assay

Apoptotic cells were visualized on paraffin-embedded tissue sections using a Click-iT TUNEL Alexa Fluor 488 kit (ThermoFisher Scientific) according to the manufacturer’s instructions.

#### Image acquisition and processing

Immunofluorescence images were acquired using a Leica TCS SP8 confocal fluorescence microscope (Leica Microsystems) with HC PL FLUOTAR 25×/0.75 NA and HC PL APO 63×/1.4 NA immersion oil objectives. For tissue sections, z-stacks were acquired, and representative maximal projections were shown. Super-resolution microscopy was performed using either a DeltaVision OMX microscope with a Blaze SIM module (GE Healthcare Life Sciences) and an U APO N 100×/1.49 NA immersion oil objective (K8 staining) or a Leica TCS SP8 STED 3X microscope (Leica Microsystems) with an HC PL APO 100×/1.4 NA immersion oil objective (actin and tubulin staining). Raw images were deconvolved using Huygens Essential 4.0.0 software (Huygens; Scientific Volume Imaging). Representative maximal projections of z-stacks are shown. Bright-field images were acquired on a Leica DM6000 wide-field microscope with an HC PLAN APO 20×/0.7 NA dry objective. Post-acquisition processing was performed with Photoshop CS6 (Adobe Systems Inc., Mountain View, CA) and the open-source Fiji image processing package^3^.

#### Transmission electron microscopy

Immediately after a dissection, the distal colon free of feces was cut into small pieces (3×3 mm) and fixed in 2.5% glutaraldehyde in Sorensen’s buffer. Samples were post-fixed with 1% OsO_4_ in Sorensen’s buffer, contrasted with 1% uranyl acetate in 50% ethanol overnight, dehydrated through a graded ethanol series followed by propylene oxide, and embedded into an Epon 812 substitute and Durkupan ACM (Sigma-Aldrich). Polymerized blocks were cut into 80-nm-thin sections, contrasted with an aqueous solution of uranyl acetate and inspected using a Morgagni 268 transmission electron microscope operated at 80 kV. Images were captured using a Mega View III CCD camera (Olympus Soft Imaging Solutions).

#### Histological and morphometric analyses

Blinded histopathology evaluation of human colon biopsy samples stained with H&E was independently performed by two trained pathologist (E.S. and L.B.). Blinded histopathology evaluation of mouse colon sections stained with H&E and PAS was performed by a trained pathologist (J.S.). The numbers of Ki-67-, PAS-, ChgA-, K20-, and TUNEL-positive cells, and the numbers of detached cells per crypt (colon) or villus (small intestine) were counted and normalized to the total numbers of IECs. At least 7 crypts or villi were analyzed per mouse and genotype. Crypt and villus damage was assessed as a percentage of crypts/villi with >10% of IECs detached from the BM. To analyze migration of IECs in the colonic crypt, the relative position of each BrdU-positive cell within the crypt was assessed. For scoring the cell position, cells were numbered sequentially from the crypt base to lumen, with cell position 0 being occupied by the first cell at the base of each crypt. The Muc2-positive area was measured and quantified using Fiji software. At least 8 images were analyzed per genotype.

#### In vivo intestinal permeability assay

To measure intestinal permeability, FITC-dextran 4 (4000 MW; TdB Consultancy) dissolved in PBS was administered by oral gavage (0.6 g/kg body weight) to mice after a 4-hour fast. Blood was obtained by retro-orbital bleeding from anesthetized mice 4 hours later and collected in heparin-coated tubes (Microvette® CB 300, Sarstedt). Then plasma was separated. Serum-FITC levels were measured at 488 nm using an Envision 2104 MultiLabel Reader (PerkinElmer).

#### Ex vivo intestinal transepithelial electrical resistance measurement

The Ussing chamber technique was used to measure intestinal transepithelial electrical resistance measurement (TEER). Whole-thickness segments of the proximal and distal colon were mounted in Ussing chambers (exposed area 0.096 cm^2^) filled with a Krebs-Ringer solution containing (in mM) Na^+^ (140.5), K^+^ (5.4), Ca^2+^ (1.2), Mg^2+^ (1.2), Cl^−^ (123.8), HCO_3_ (21), HPO_4_ (2.4), H_2_PO_4_ (0.6), glucose (10), mannose (10), glutamine (2.5), and β- hydroxybutyrate (0.5). The segments were permanently oxygenated with a mixture of 95% oxygen and 5% carbon dioxide (pH 7.4, 37 °C). After 30 min of equilibration, TEER was measured using bipolar rectangular current pulses (10 μA, 200 ms) and a programmable voltage-clamp device (Müssler Scientific Instruments).

#### In vitro myeloperoxidase activity measurement

The distal colon was cut into small pieces and homogenized in a 50 mM phosphate buffer, pH 6, with 0.5% cetrimonium bromide (50 mg tissue/ml of buffer) and incubated at 60°C for 2 hours. Myeloperoxidase (MPO) activity was assessed in a clear supernatant using 3,3’,5,5’-tetramethylbenzidine (Sigma-Aldrich) as a substrate as described before^4^. Final activity is expressed in U per mg of protein.

#### Protein extraction and immunoblotting

Excised proximal and distal colons or ilea were cut open longitudinally and washed with PBS on ice. The colonic mucosa was scraped off using square coverslips. Snap-frozen mucosal scrapings were homogenized in ice-cold RIPA (20 mM Tris-HCl pH 7.5, 150 mM NaCl, 1 mM Na_2_EDTA, 1 mM EGTA, 1% NP-40, 0.5% SDS supplemented with Halt protease and a phosphatase inhibitor Cocktail (Thermo Fisher Scientific)) using the Tissue Lyzer II (Qiagen). Caco-2 cells were lysed in ice-cold RIPA by shearing through a 29G needle. Protein concentrations were determined using a BCA Protein Assay Kit (Thermo Fisher Scientific). Clarified lysates were resolved on SDS-PAGE and transferred to a nitrocellulose membrane for immunodetection. The following primary antibodies were used: ZO-1 (61-7300, ThermoFisher), E-cadherin (610181, BD Biosciences), desmoglein (611002, Progen), integrin α6 (ab181551, Abcam), integrin β4 (ab182120, Abcam), GAPDH (G9545, Sigma-Aldrich), keratin 8 (Troma I, Developmental Studies Hybridoma Bank), keratin 19 (Troma III, Developmental Studies Hybridoma Bank), and keratin 18 (Ks18.04, Progen). The following secondary antibodies were used: HRP-conjugated goat anti-guinea pig IgG (Sigma-Aldrich), donkey anti-mouse IgG (IRDye 680RD), donkey anti-rabbit IgG (IRDye 800CW), and goat anti-rat (IgG IRDye 800CW; all Licor). Signals were detected with an ECL Plus Western Blotting Detection System (GE Healthcare Life Sciences) and recorded with a Luminescent Image Analyzer LAS-3000 (Fujifilm Life Science, Düsseldorf, Germany) or the Odyssey 9120 imaging system (Licor). The densitometry of blots was analyzed using QuantiScan version 1.5 software (Biosoft).

#### Quantitative reverse transcriptase PCR

RNA was isolated from snap-frozen mucosal scrapings (see above) using TRI reagent (Sigma-Aldrich) according to the manufacturer’s instructions. The RNA concentration was determined using the Nanodrop ND-1000 (Thermo Fisher Scientific). cDNA was prepared using M-MLV reverse transcriptase (ThermoFisher Scientific) with random oligo(dT)18 primers. qPCR was performed with SYBR Green JumpStart Taq ReadyMix (Sigma-Aldrich) using gene-specific primers (Supplementary Table 2). Due to previously reported instability of reference genes in colitic mice^5^, expression of several reference genes was compared in *Ple^fl/fl^* and *Ple^ΔIEC^* mice (Actb, Eef2, GAPDH, Hmbs and Tbp). As expression of none of these genes significantly differed between *Ple^fl/fl^* and *Ple^ΔIEC^*, relative RNA expression was calculated by the comparative threshold cycle method (ΔΔCt) using a GAPDH internal reference gene control.

#### Cell stretching

Stretch experiments were carried out on flexible polydimethylsiloxane (PDMS, Sylgard) substrates with 4.0 cm^2^ internal surface. The stretcher had a linear stage for a uniaxial stretch and was driven by a computer-controlled stepper motor ^6^. The substrates were coated with 50 μg/ml laminin or collagen type I in PBS at 4°C overnight, and 50,000 cells were seeded 24 hours prior to experiments. A uniaxial cyclic stretch was performed in an incubator under normal cell culture conditions (37°C, 5% CO_2_, 95% humidity) for 1 hour at 10, 20, 30 and 50% stretch amplitude (peak-to-peak).

#### Radial shear assay

Radial shear assay was performed on a customized spinning disk device^7^ consisting of a rotating glass plate driven by compressed air. The glass plate was located approximately 300 μm above a 35 mm plastic dish with adherent cells seeded at a density of 15000 cells/dish. The shear force was generated by a rotational speed of 1500 rpm and applied for 5 min. To assess the cell density, images of areas defined by radial distances 2-4 mm of the dish (corresponding to 0.7-1.5 Pa shear stress) were acquired before and after spinning. Then cells were stained with propidium iodide (PI; Fischer Scientific), and fractions of dead (PI-positive) and detached cells were calculated.

#### Magnetic tweezer microrheology

To determine the strength of cell-matrix adhesion, 5.09 μm carboxylated super-paramagnetic beads (microParticles GmbH) were coated with 20 μg/100 μl laminin or collagen type I (in PBS). The bead slurry (50%) was added to Caco2 and hCC cells grown on a 35 mm plastic dish and incubated for 1 hour under standard conditions. A magnetic field was generated as previously described^6^ using a solenoid with a needle-shaped core (HyMu80 alloy, Carpenter). The needle tip was placed at a distance of 20 µm from a bead bound to the cell surface using a motorized micromanipulator (Injectman NI-2, Eppendorf). During measurements, bright-field images were taken by a CCD camera (ORCA ER, Hamamatsu) at a rate of 40 frames/s. The median of bead detachment (50% of adherent beads), determined under increasing forces of up to 15 nN, was used for calculating cumulative rupture forces.

#### Faecal microbiota analysis

Stool samples were collected from 6 Ple^fl/fl^ and 6 *Ple^ΔIEC^* mice at the age of 4, 12 and 20 weeks and analyzed for bacterial composition as described earlier^8^. Briefly, genomic DNA was extracted with a MasterPure™Complete DNA and RNA Purification Kit (Epicentre) with repeated bead-beating in Lysing Matrix Y tubes using a FastPrep homogenizer (both MP Biomedicals). Next, the V3-V4 region of the 16S rRNA gene was amplified using barcoded bacterial 16SrRNA-specific primers 341F (5’-CCTACGGGNGGCWGCAG-3’) and 806R (5’-GGACTACHVGGGTWTCTAAT-3’). PCR amplification was performed with KAPA 2G Robust Hot Start DNA Polymerase (Kapa Biosystems), with following concentrations: Buffer B 1×, Enhancer 1×, dNTP 0.2 mM each, primers 0.5 μM each, DNA sample 4 ng/μl, KAPA polymerase 0.5 U. Cycle parameters were 3 min 94°C, 25 cycles of 30 s at 94°C, 1 min at 54.2°C, and 1 min 15 s at 72°C; the final extension was at 72°C for 10 min. Three PCR products were pooled to minimize random PCR bias, and the length of PCR products was checked by agarose gel electrophoresis. Equal amounts of each sample were plate-purified using the SequalPrep™Normalization Plate (96) Kit (Invitrogen). Then equimolar amounts of PCR products from each sample were pooled, and MiSeq platform compatible adapters were ligated using a TruSeq DNA PCR-Free LT Kit (Illumina). The libraries were quantified using a KAPA Library Quantification Kit (Illumina) and sequenced on a MiSeq platform using a 2× 300bp kit at the CEITEC Genomics Core Facility.

Sequencing data were processed using QIIME (Quantitative Insights Into Microbial Ecology) version 1.9.1^9^. Quality filtering, chimera detection, read demultiplexing, and read clustering were done as described previously^10^. Raw reads were demultiplexed and quality filtered, and all sequences containing unknown base calls were excluded. Chimeric reads were detected and discarded using USEARCH algorithms^11^. The final dataset contained 80,185 high-quality reads (median 1,306 reads per sample, range 28 - 8766). To make samples comparable, 4 samples with fewer than 460 reads were removed from the analysis, and the OUT table was rarefied at a depth of 460 sequences per sample. Operational taxonomic units (OTUs) were clustered at a 97% similarity level, and representative sequences were identified using a Ribosomal Database Project classifier^12^ against bacterial GreenGenes database 13.8^13^. For a microbiota analysis, PD whole tree metrics measuring the total descending branch length in the phylogenetic tree for each OTU was used to describe alpha diversity. The Principle Coordinate Analysis (PCoA) based on unweighted UniFrac distance metrics was used to describe beta diversity.

The sequence data are available in the Sequence Read Archive (SRA; http://www.ncbi.nlm.nih.gov/sra) under BioProject accession number PRJNA561691.

#### High salt extraction of Caco-2 cells

High salt extraction of Caco-2 cells was performed as described previously^14^. Cell fractions were prepared by solubilizing cells for 2 min at 4°C with a buffer containing 1% TX-100, 5 mM EDTA, and Halt Protease Inhibitor Cocktail (Thermo Fisher Scientific) in PBS pH 7.4, followed by centrifugation (16,000 × g, 10 min). The supernatant was collected as a soluble fraction. The pellet was homogenized in 1 ml of 10 mM Tris-HCl pH 7.6, 140 mM NaCl, 1.5 M KCl, 5 mM EDTA, 0.5% Triton X-100, supplemented with Halt Protease Inhibitor Cocktail. After 30 min (at 4°C), the homogenate was pelleted (16,000×g; 10 min), and the pellet (insoluble fraction) was rehomogenized with 5 mM EDTA in PBS pH 7.4. The resulting homogenate was further centrifuged (16,000×g; 10 min) to obtain the insoluble keratin-enriched high salt extract (HSE). All fractions were resolved by SDS-PAGE and their composition was analyzed by immunoblotting.

### 2. Supplementary FIGURE LEGENDS

**Figure S1.**
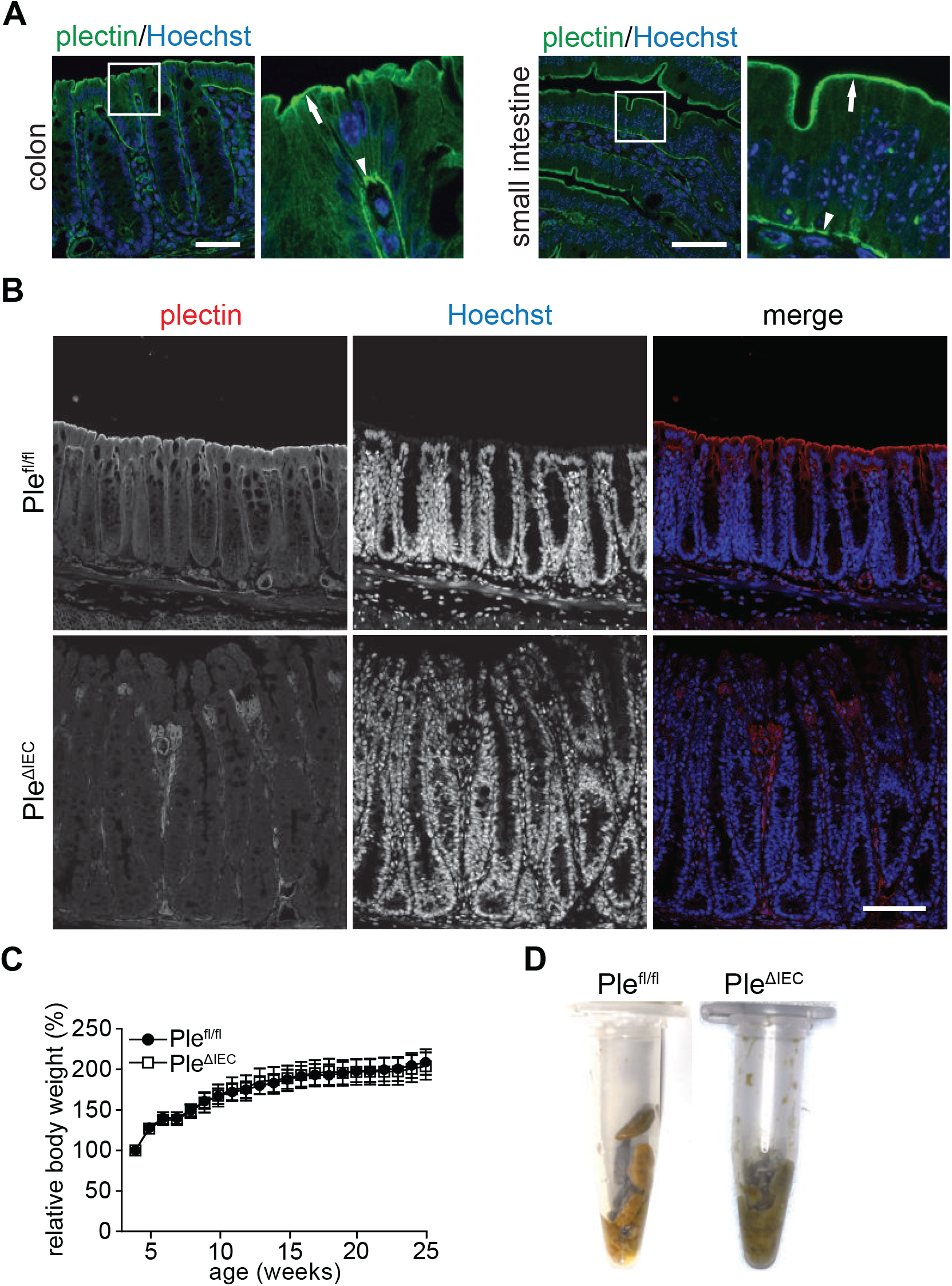
Immunolocalization of plectin in mouse intestine and phenotypic alterations of colonic epithelium, body weight, and stool consistency in plectin-deficient (*Ple^ΔIEC^*) mice. (A) Paraffin-embedded sections from distal colon and small intestine of *Ple^fl/fl^* mice were immunolabelled with antibodies to plectin (green). Nuclei were stained with Hoechst (blue). Plectin staining at the apical (arrows) and basal (arrowheads) membranes,. Scale bar, 50 μm. Boxed areas show ×4.5 images. (B) Paraffin-embedded distal colon sections from *Ple^fl/fl^* and *Ple^ΔIEC^* mice were immunolabelled with antibodies to plectin (red). Nuclei were stained with Hoechst (blue). Scale bar, 100 μm. Note immunofluorescence signal in plectin-positive mesenchymal niche of both *Ple^fl/fl^* and *Ple^ΔIEC^* colons. (C) Body weights of *Ple^fl/fl^* and *Ple^ΔIEC^* mice were followed for 25 weeks. Graph shows relative body weight normalized to birth body weight. n = 7. (D) Stool consistency in *Ple^fl/fl^* and *Ple^ΔIEC^* mice.

**Figure S2.**
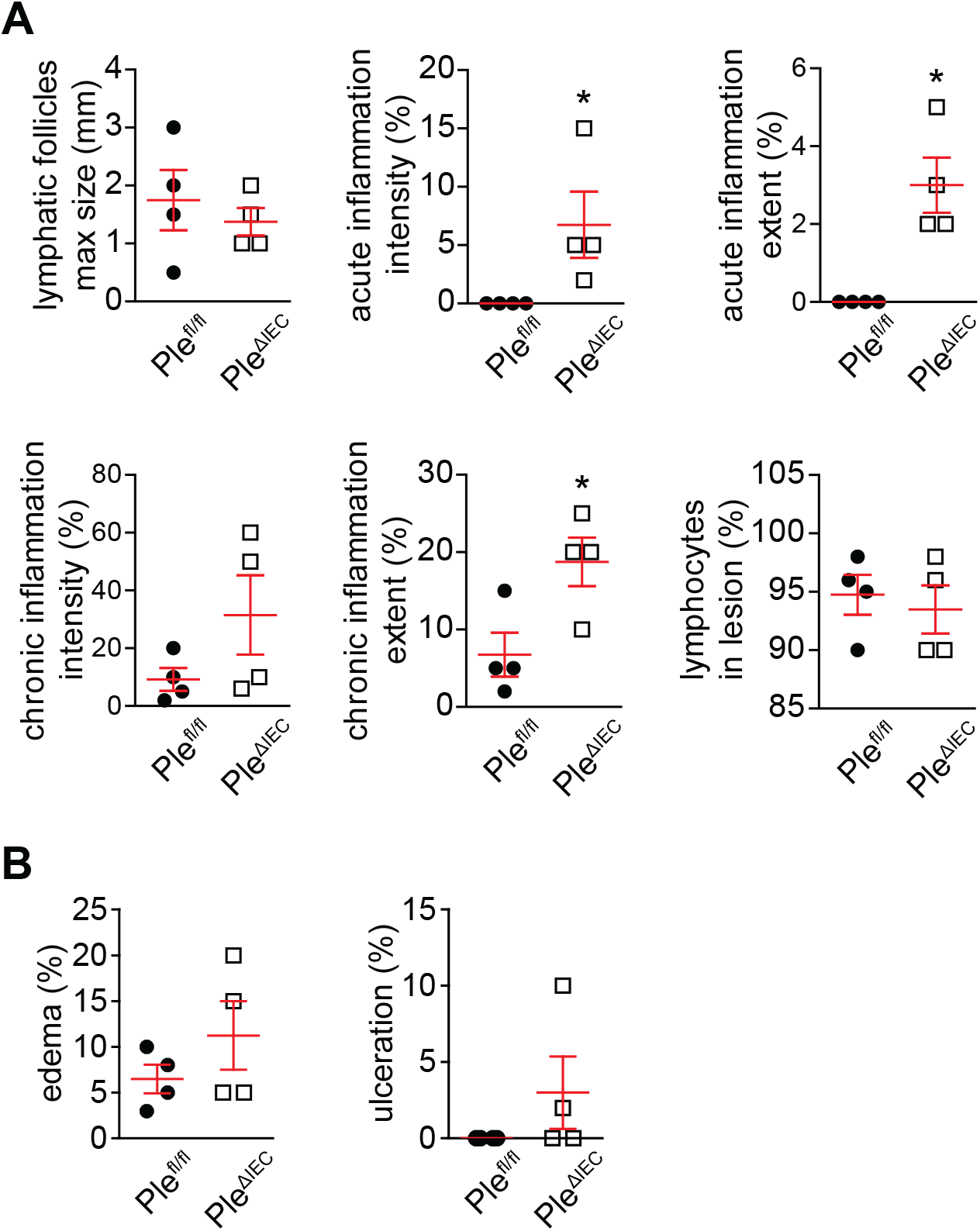
Histological assessment of colonic tissue damage and inflammation in *Ple^ΔIEC^* compared to *Ple^fl/fl^* mice. (A) Quantification of inflammatory parameters on H&E-stained sections of *Ple^fl/fl^* and *Ple^ΔIEC^* colons. Graphs show maximal (max) sizes of lymphatic follicles, percentage of acute inflammation intensity, acute inflammation extent, chronic inflammation intensity, chronic inflammation extent, and lymphocytes in lesion. (B) Quantification of tissue damage assessed from H&E-stained sections of *Ple^fl/fl^* and *Ple^ΔIEC^* colons (percentage of edema and ulceration). n = 4. Data are presented as mean ± SEM, \**P* < 0.05.

**Figure S3.**
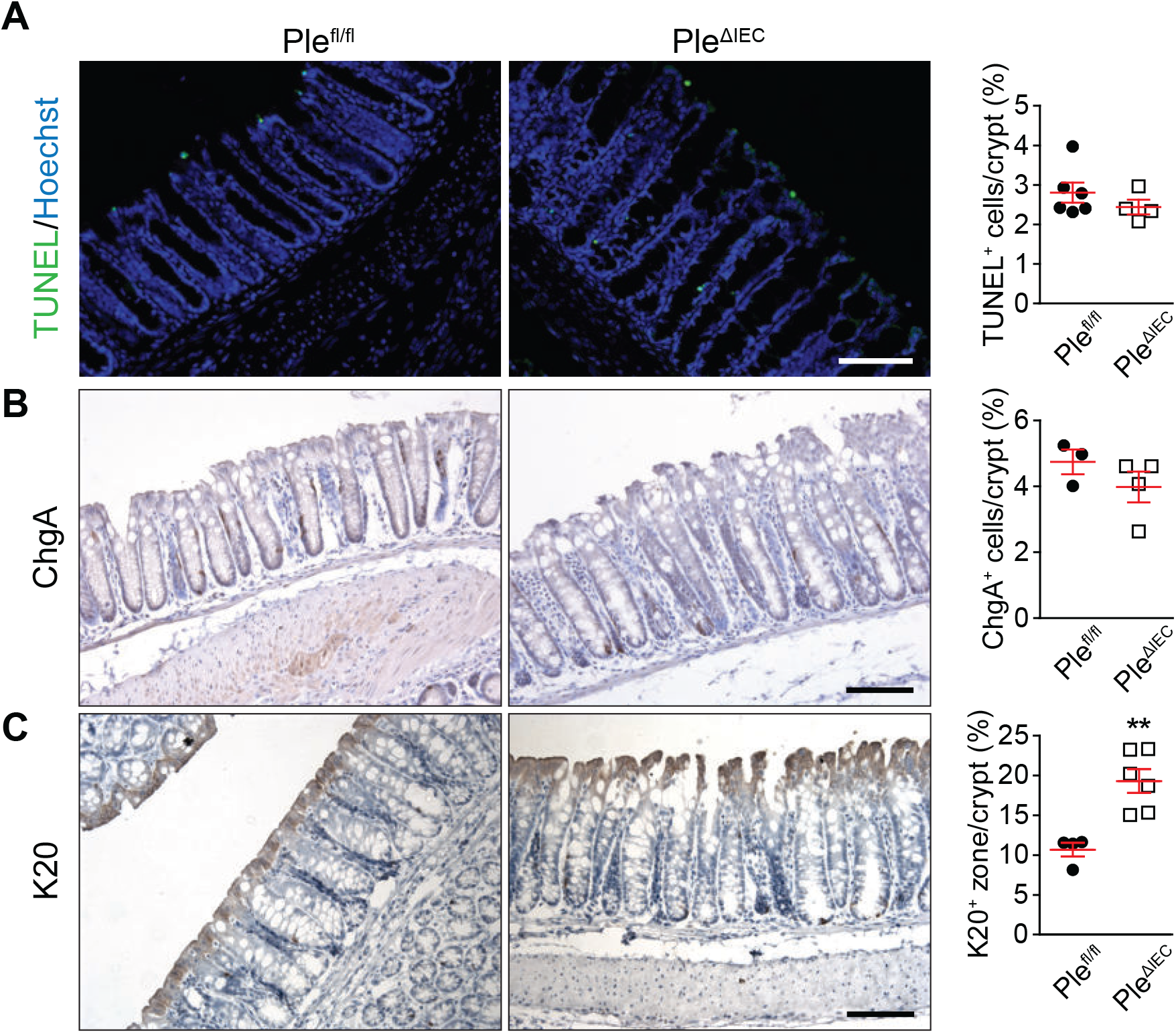
Characterization of colonic crypts in *Ple^fl/fl^* and *Ple^ΔIEC^* mice. (A-C) Representative images of fluorescent TUNEL staining of apoptotic cells (green) (A), ChgA immunohistochemistry (enteroendocrine cells) (B), and K20 immunohistochemistry (mature IECs) (C) of *Ple^fl/fl^* and *Ple^ΔIEC^* colon sections. Nuclei in (A) were stained with Hoechst (blue). Scale bars, 100 μm. Corresponding graphs show percentage of positively labeled (+) IECs and K20 zone per crypt cells. n = 3-6. Data are presented as mean ± SEM, \*\**P* < 0.01.

**Figure S4.**
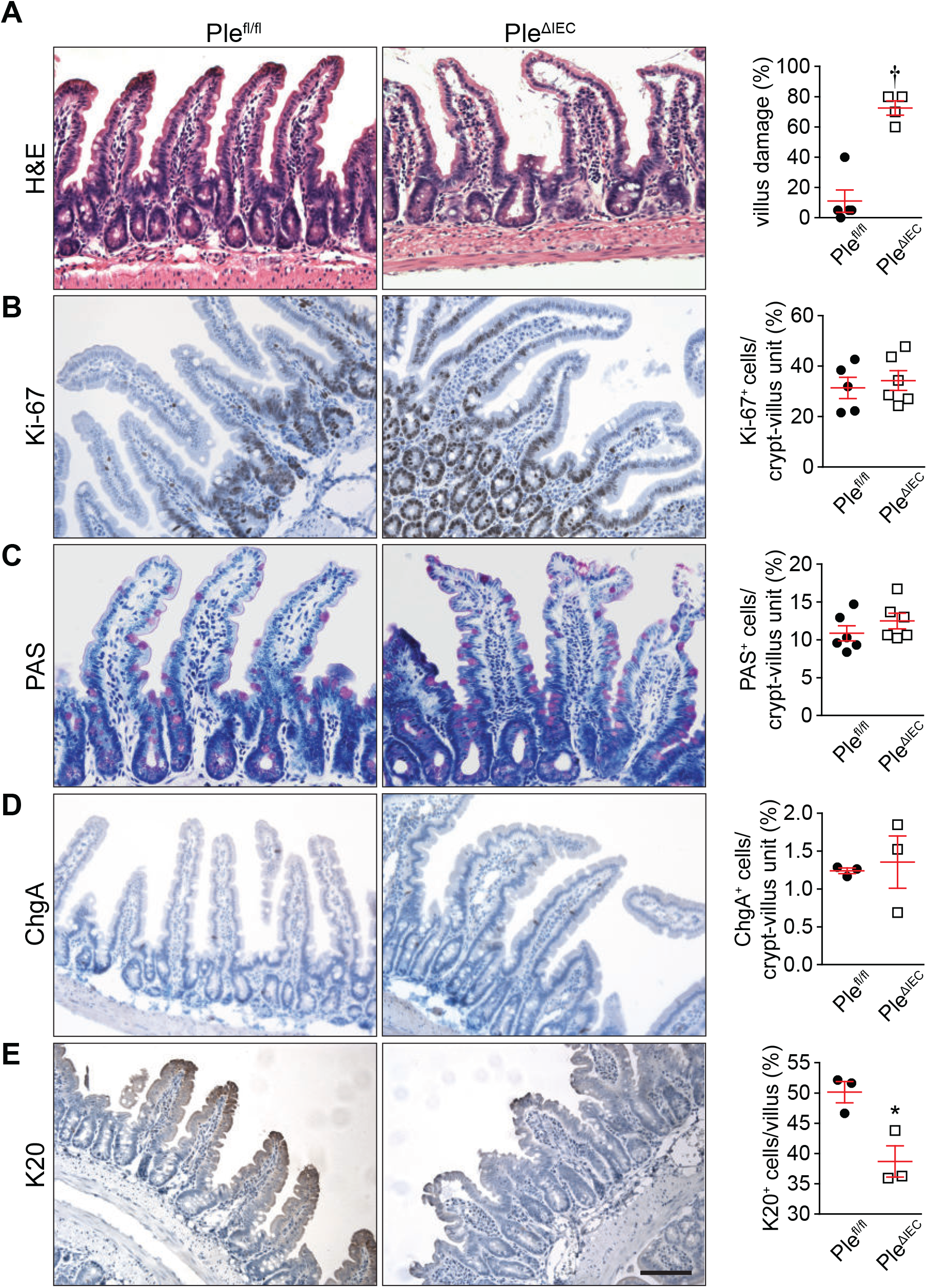
Characterization of small intestinal crypt-villus units in *Ple^fl/fl^* and *Ple^ΔIEC^* mice. (A-E) Representative images of H&E staining (A), Ki-67 immunohistochemistry (proliferating cells) (B), PAS staining (goblet cells) (C), ChgA immunohistochemistry (enteroendocrine cells) (D), and K20 immunohistochemistry (mature IECs) (E) of small intestine sections. Scale bar, 100 μm. Corresponding graphs show percentage of damaged crypt-villus units (villus damage given as percentage of villi with > 10% of IECs detached from BM)A) and percentage of the positively labeled (+) IECs per crypt-villus unit (B, C) or villus (D, E). n = 3-6. Data are presented as mean ± SEM, \**P* < 0.05, †*P* < 0.001.

**Figure S5.**
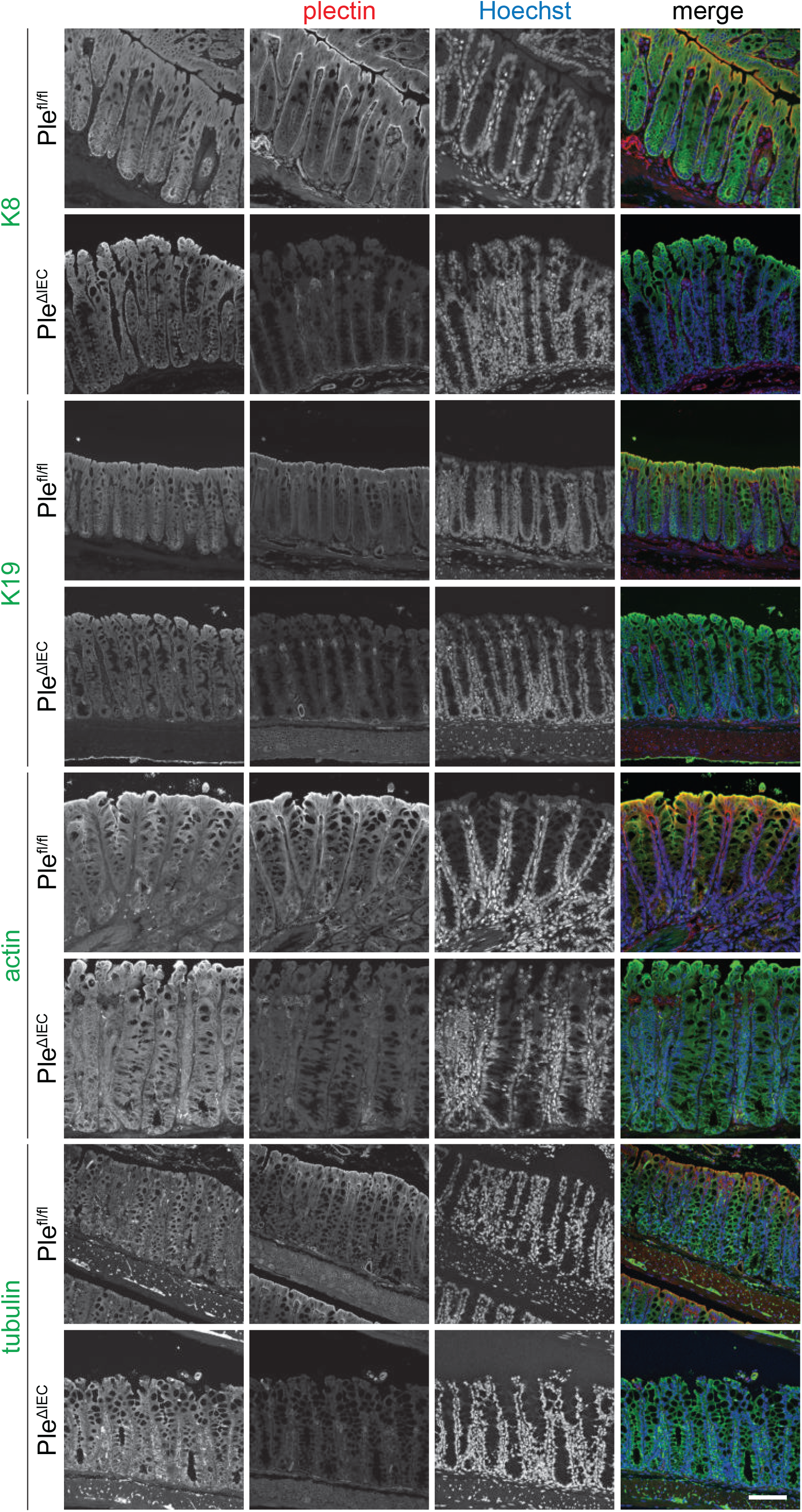
Organization of KFs, actin fibers, and microtubules in colonic epithelium of *Ple^fl/fl^* and *Ple^ΔIEC^* mice. Sections of *Ple^fl/fl^* and *Ple^ΔIEC^* distal colon were immunolabelled with antibodies to K8, K19, β-actin, tubulin (all green), and plectin (red). Nuclei were stained with Hoechst (blue). Representative overview images are shown (see Figure 5A for more detailed images of pan-K-immunolabelled colons). Scale bar, 100 μm.

**Figure S6.**
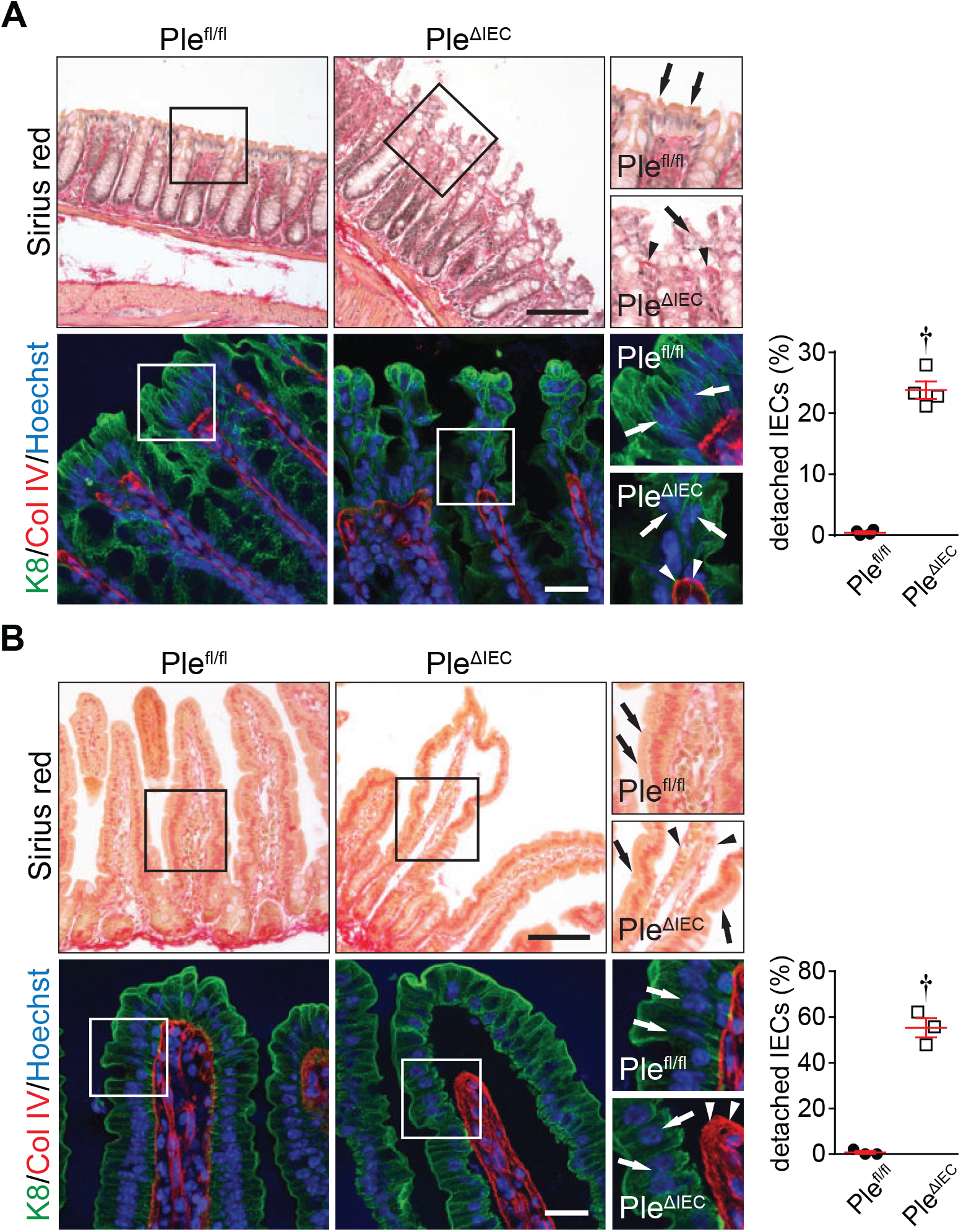
Detachment of IECs from BM in *Ple^ΔIEC^* intestinal epithelium. (A, B) Sections of *Ple^fl/fl^* and *Ple^ΔIEC^* distal colon (A) and small intestine (B) were stained for fibrillar collagen with Sirius red (upper panels) and immunolabelled for K8 (green) and Col IV (red; lower panels). Nuclei were stained with Hoechst (blue). Boxed areas show ×1.5x images. The arrows, IECs; the arrowheads, BM. Scale bars, 100 μm (upper panels) and 25 μm (lower panels). Graphs show percentage of detached IECs per crypt (A) or villus (B). Data are presented as mean ± SEM, †*P* < 0.001.

**Figure S7.**
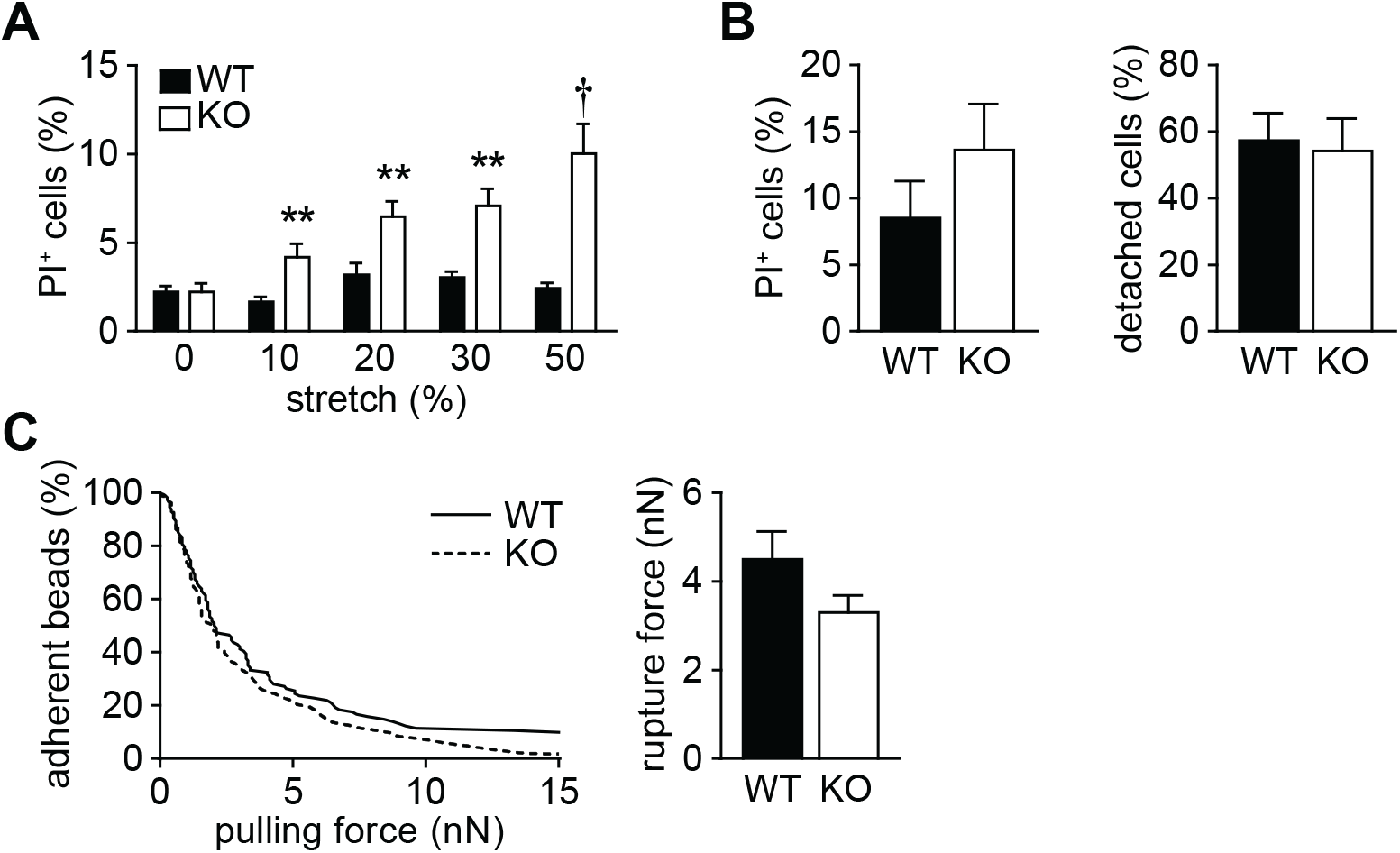
Increased mechanical vulnerability and attenuated adhesion of KO hCC cells. (A) Quantification of viability of WT and KO hCC cells exposed to uniaxial cyclic stretch shown as percentage of dead (PI^+^) cells. n = 9-12. (B) Quantification of WT and KO hCC cell viability (left) and adhesion (right) under radial shear flow shown as percentage of dead and detached cells, respectively. n = 8. (C) Adhesion strength between ECM-coated paramagnetic beads and WT and KO hCC cells adhesions was quantified using magnetic tweezers. Left graph shows bead detachment (percentage of adherent beads at given pulling force); right graph shows cumulative rupture force (calculated from median bead detachment). n = 103. Bar graphs represent mean ± SEM, \*\**P* < 0.01, †*P* < 0.001.

**Figure S8.**
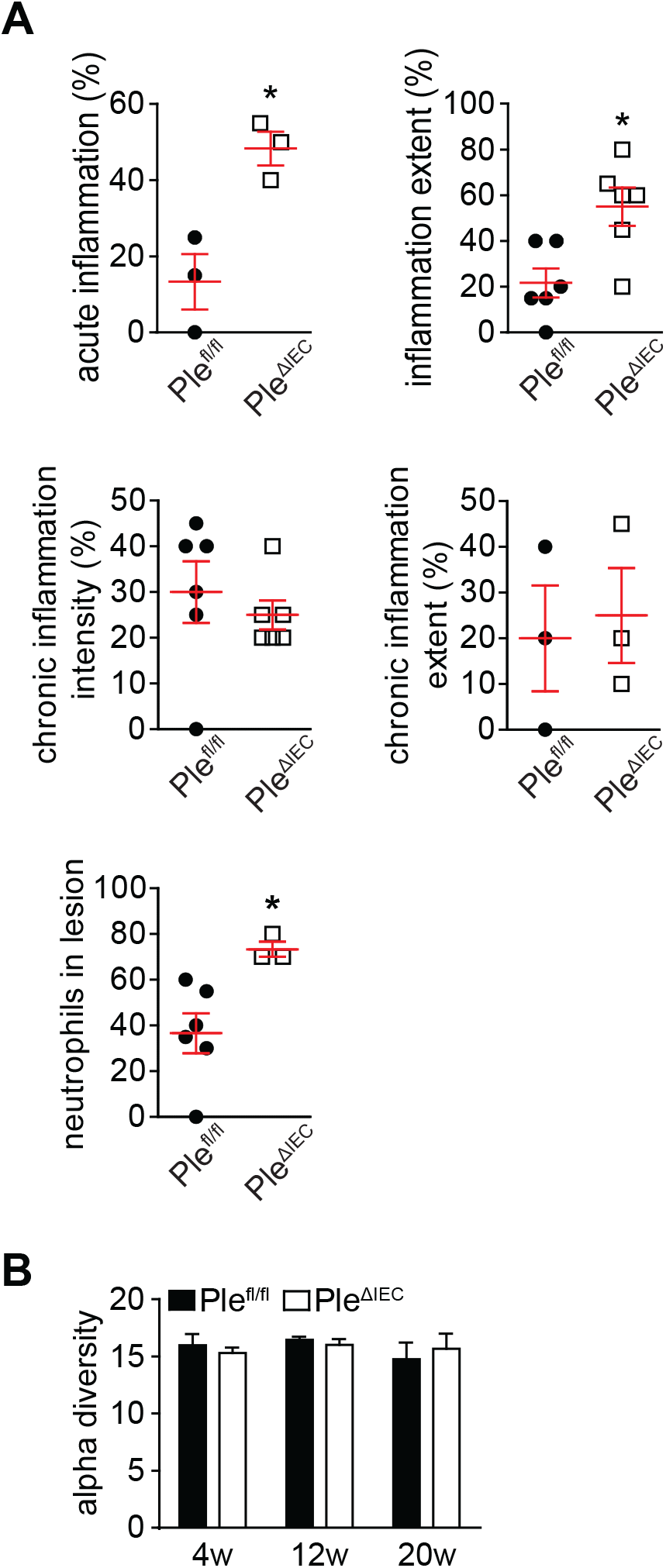
DSS-induced colitis in *Ple^ΔIEC^* and *Ple^fl/fl^* mice. (A) Quantification of inflammatory parameters on H&E-stained colonic sections from DSS-treated *Ple^fl/fl^* and *Ple^ΔIEC^* mice. Graphs show percentage of acute inflammation, inflammation extent, chronic inflammation intensity, chronic inflammation extent, and the number of neutrophils in lesion. n = 3-6.. (B) Alpha diversity of fecal microbiota in 4-, 12-, and 20-week-old untreated *Ple^fl/fl^* and *Ple^ΔIEC^* mice determined by 16S rDNA sequencing using PD whole tree metrics. n = 4-6. Data are presented as mean ± SEM, \**P* < 0.05.

**Figure S9.**
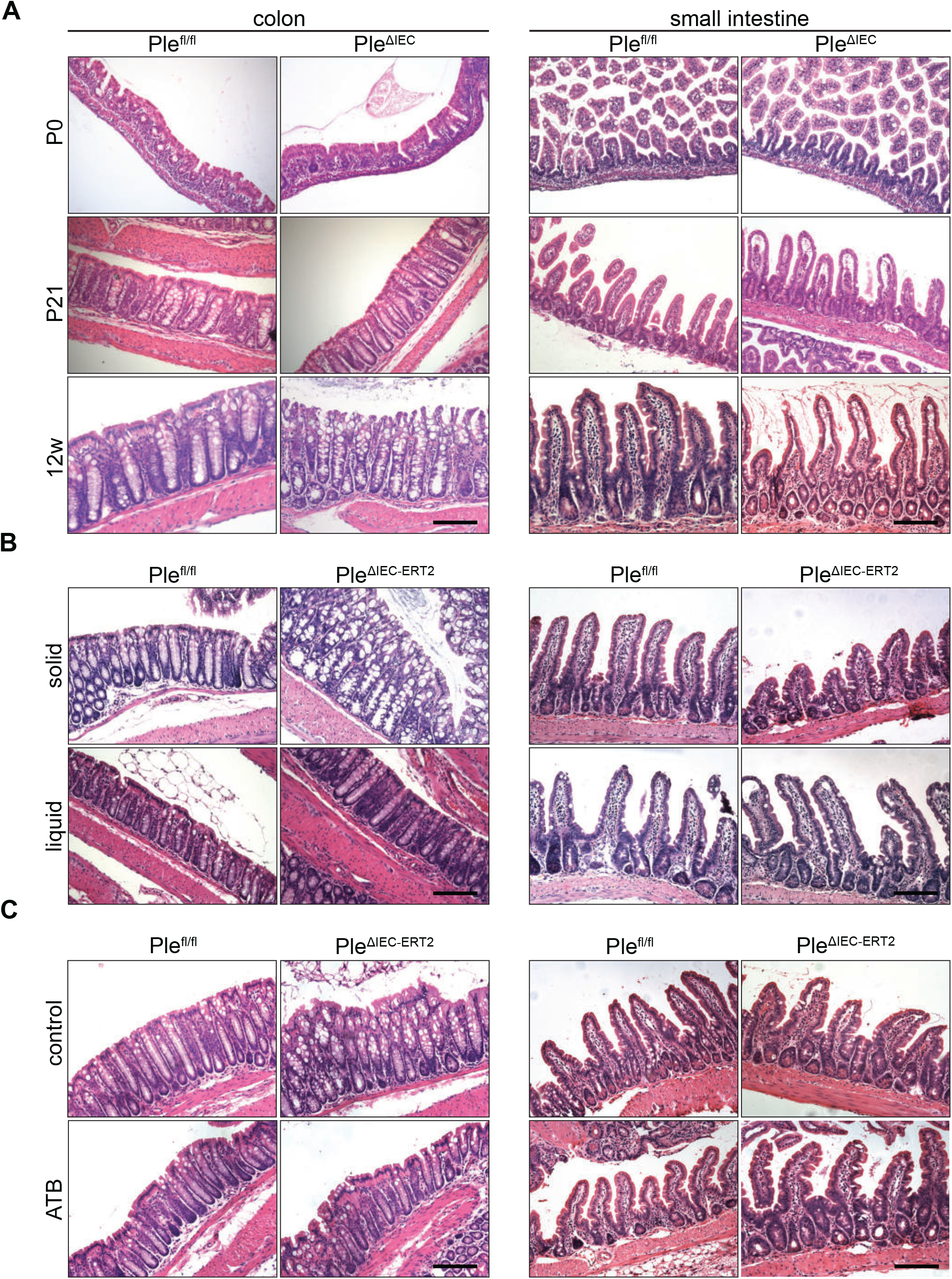
Tissue damage in colon and small intestine of WT and plectin-deficient mice at weaning, on liquid diet, and under antibiotic treatment. (A) Representative images of H&E-stained colon (left panels) and small intestine (right panels) dissected from *Ple^fl/fl^* and *Ple^ΔIEC^* mice at postnatal day 0 (P0), postnatal day 21 (P21), and at 12 weeks (12w). (B,C) Representative images of H&E-stained colon (left panels) and small intestine (right panels) dissected from 9-week-old *Ple^fl/fl^* and *Ple^ΔIEC-ERT2^* mice, either kept on solid chow or provided with liquid diet (B) or exposed to antibiotic treatment (ATB) with broad-spectrum antibiotics (C) for 14 days. In all cases, plectin inactivation was induced by 3 consecutive applications of TMX on days 6, 8, and 10 and sacrificed on day 14. Scale bar, 100 μm.

**Figure S10.**
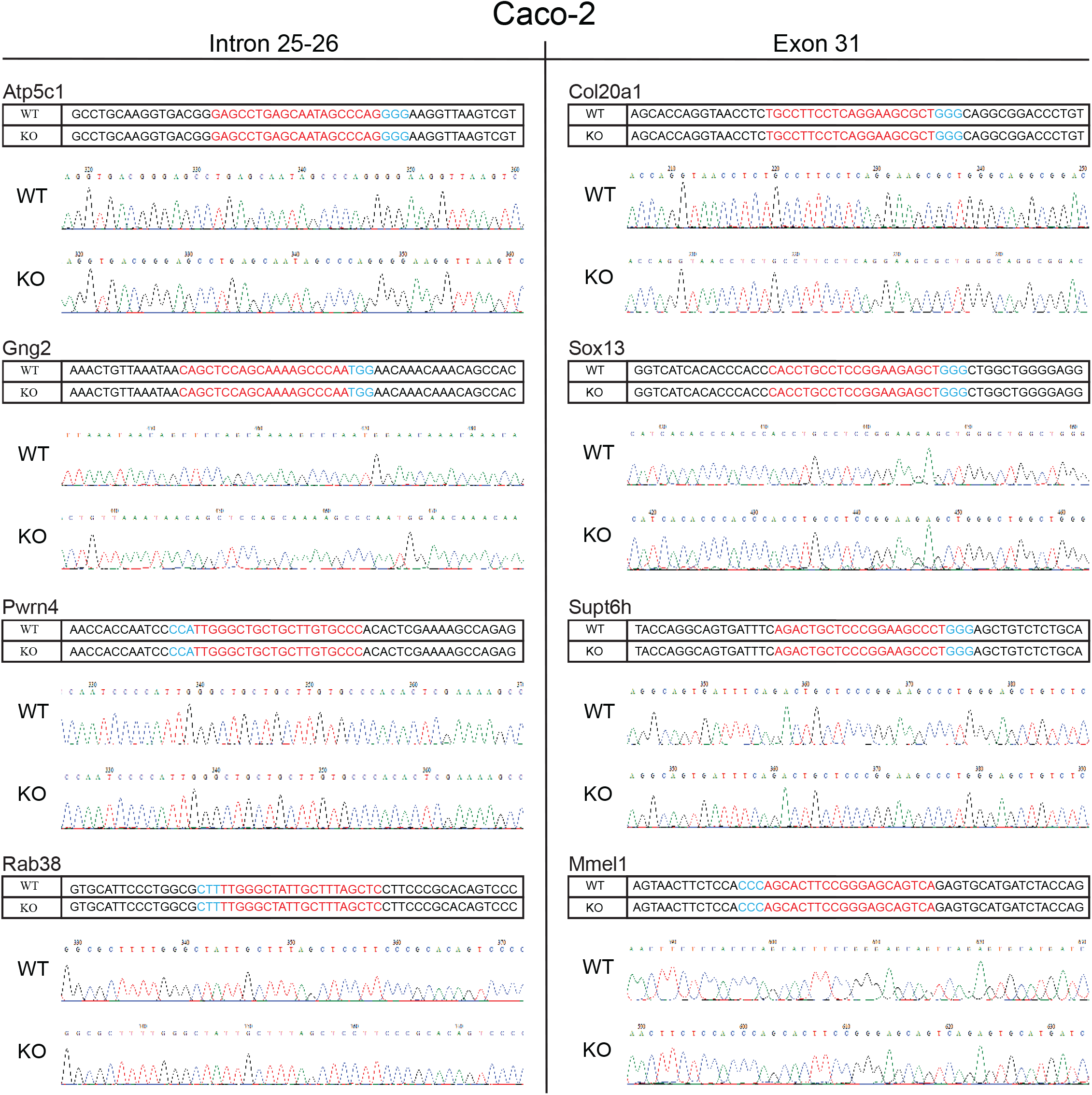
Analysis of potential off-target loci in Caco-2 cells. Four potential off-target loci for intron 25-26 (left panel) and exon 31 (right panel) guide RNA (gRNA) were amplified by PCR from genomic DNA of WT and plectin KO Caco-2 cells using gene-specific primers (listed in Supplemental Table 3). PCR products were analyzed by direct sequencing. Sequencing electropherograms and aligned sequences are shown. gRNA sequences in red; PAM sequences in blue.

**Figure S11.**
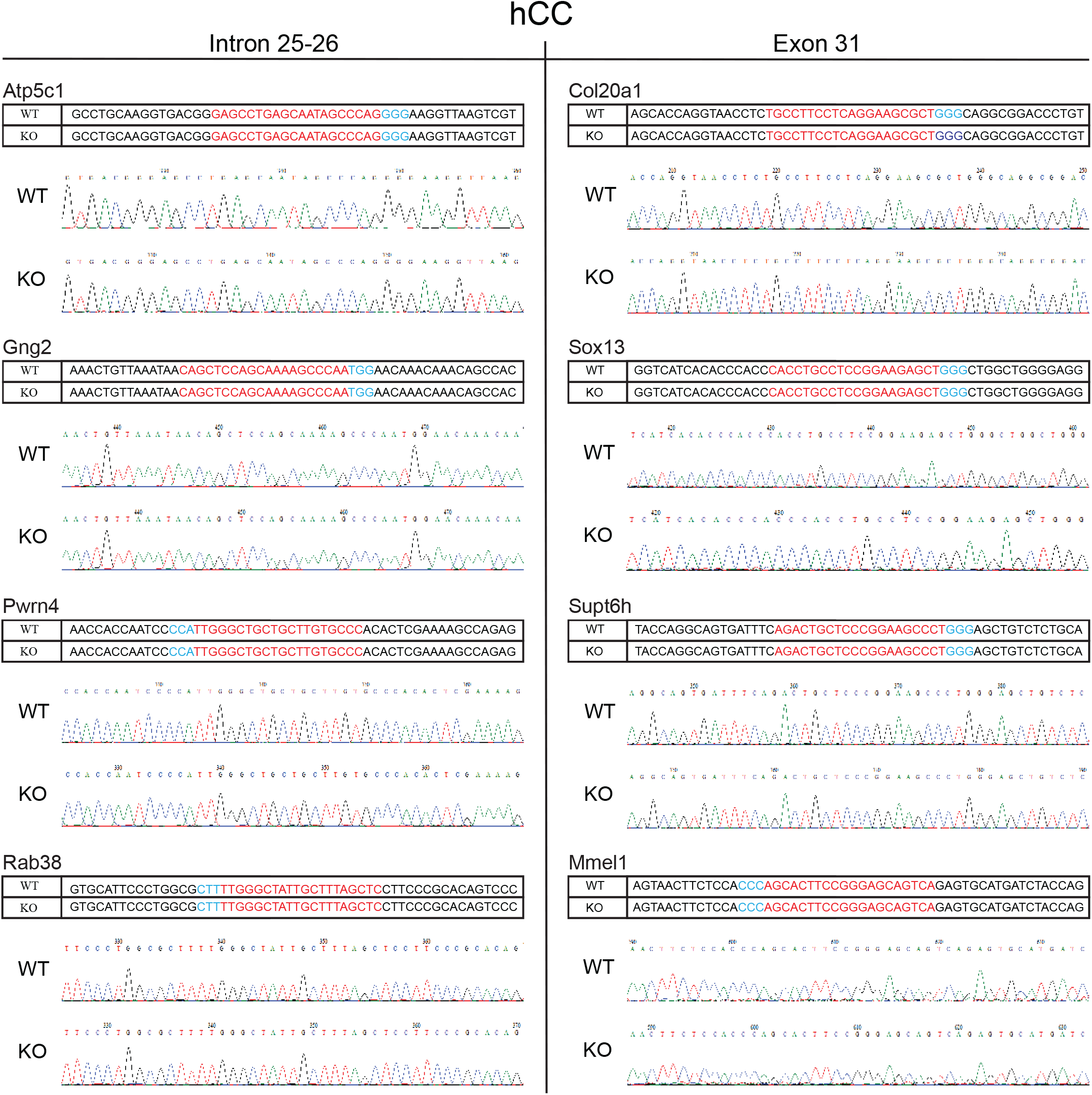
Analysis of potential off-target loci in hCC cells. Four potential off-target loci for intron 25-26 (left panel) and exon 31 (right panel) guide RNA (gRNA) were amplified by PCR from genomic DNA of WT and plectin KO hCC cells using gene-specific primers (listed in Supplemental Table 3). PCR products were analyzed by direct sequencing. Sequencing electropherograms and aligned sequences are shown. gRNA sequences in red; PAM sequences in blue.

### 3. Supplementary TABLES

**Supplemental Table 1.** Clinical and demographic descriptions of patients with ulcerative colitis (UC) and healthy patients (healthy).

|  | N | Age*<br>(years) | Gender | Duration *<br>(years) | Extension | Treatment | Mayo<sup>†</sup><br>0/1/2/3 | p-ANCA<sup>††</sup> | Inflammation<sup>†††</sup><br>0/1/2/3 |
| --- | --- | --- | --- | --- | --- | --- | --- | --- | --- |
| healthy | 18 | 47.89 (25-71) | 10/8 | - | - | - | - | - | - |
| UC | 51 | 46.57 (26-76) | 29/22 | 12.67 (1-37) | 1/14/36 | 0/22/12/17 | 19/16/10/3 | 19 | 30/5/10/6 |

N: number of patients included in the analysis
\*Mean (range)
Gender: Male/Female
Extension (UC): proctitis/left-sided/pancolitis
Treatment: None/Aminosalicylates (ASA)/Aminosalicylates + Azathioprine (ASA+AZA)/Biological treatment (Infliximab, Vedolizumab)
<sup>†</sup>Mayo referred to colonic segment used for the analysis
<sup>††</sup>p-ANCA: anti-neutrophil cytoplasm antibodies
<sup>†††</sup>Inflammation referred to colonic segment used for the analysis

**Supplemental Table 2.**
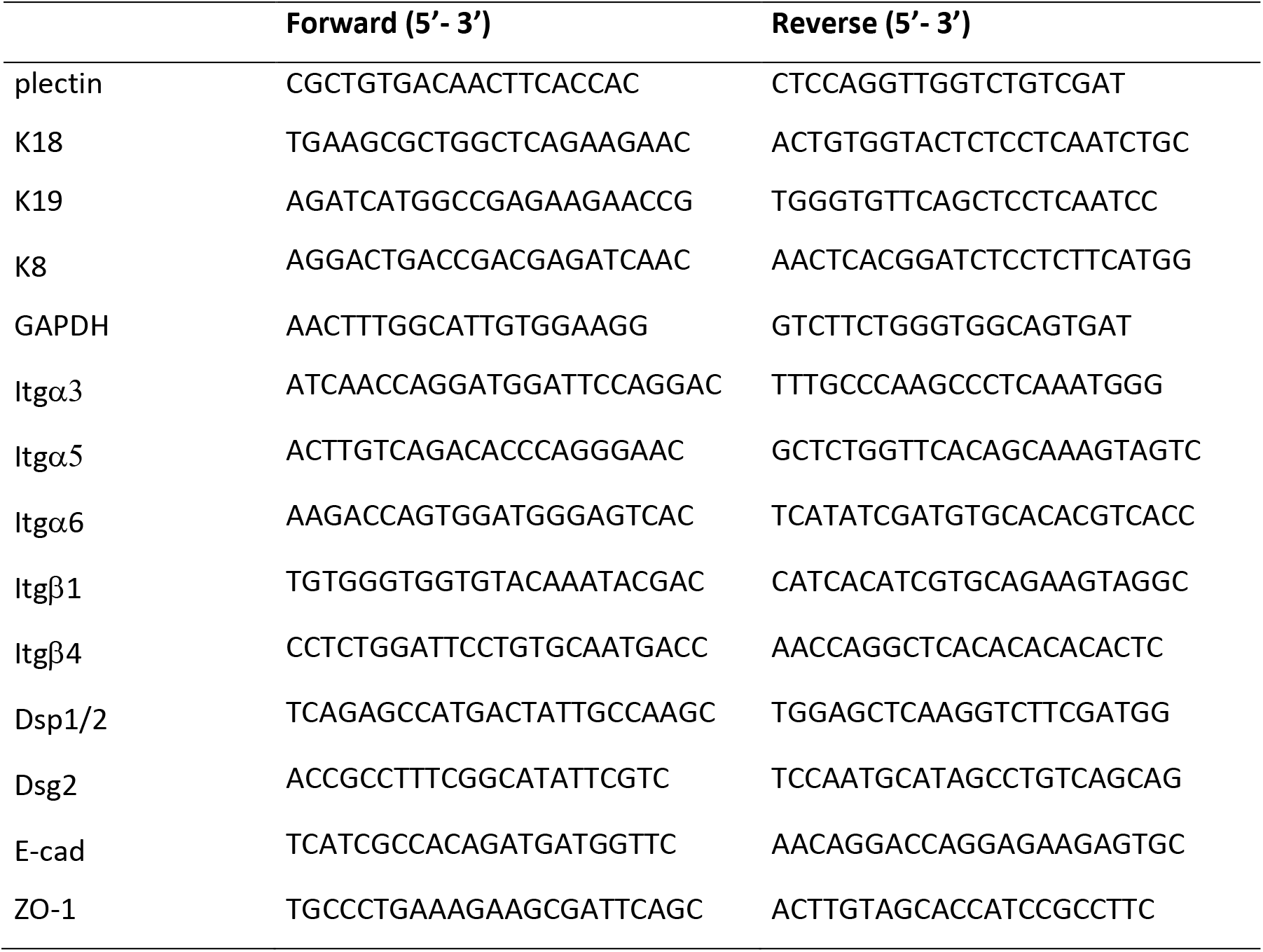
List of used RT-qPCR primers and their corresponding sequences.

**Supplemental Table 3.**
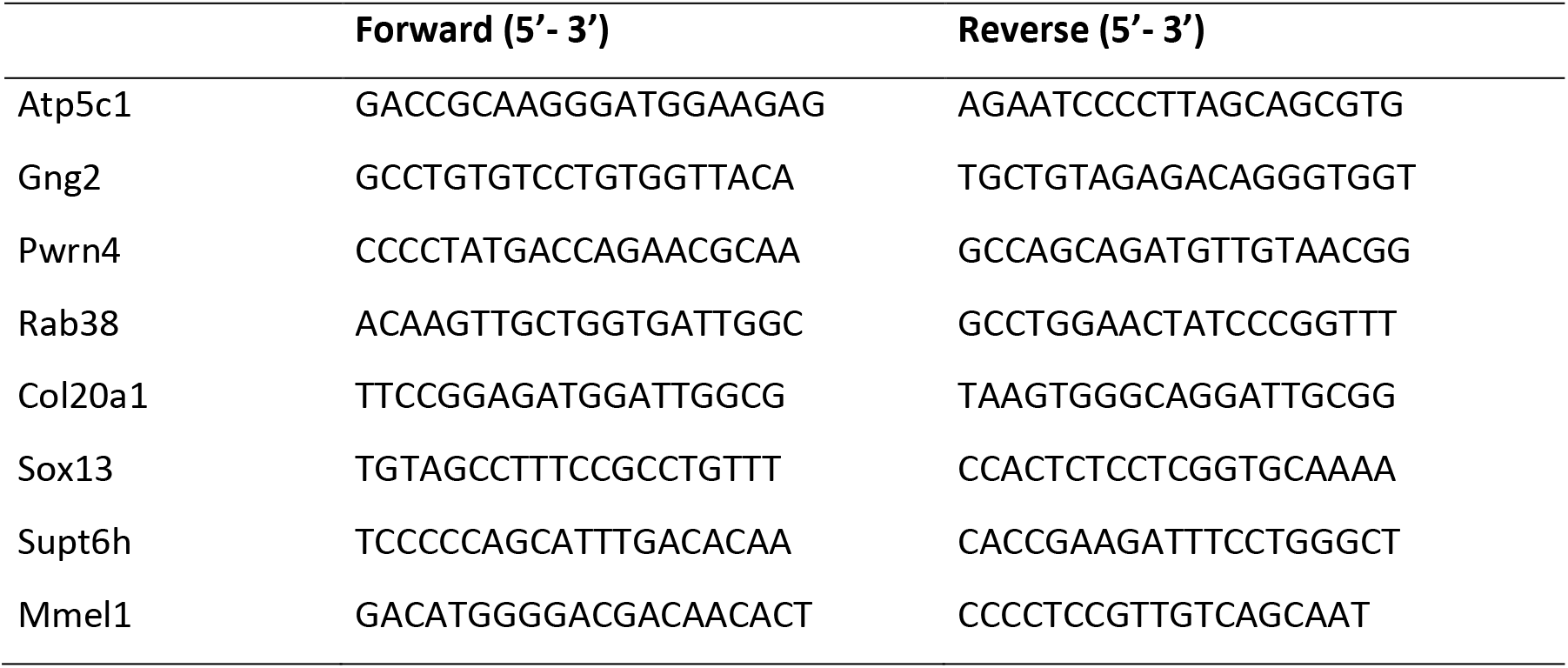
List of potential off-target genes and PCR primers with their corresponding sequences.

